# Restriction of Arginine Induces Antibiotic Tolerance in *Staphylococcus aureus*

**DOI:** 10.1101/2023.10.12.561972

**Authors:** Jeffrey A. Freiberg, Valeria M. Reyes Ruiz, Erin R. Green, Eric P. Skaar

## Abstract

*Staphylococcus aureus* is responsible for a substantial number of invasive infections globally each year. These infections are problematic because they are frequently recalcitrant to antibiotic treatment, particularly when they are caused by Methicillin-Resistant *Staphylococcus aureus* (MRSA). Antibiotic tolerance, the ability for bacteria to persist despite normally lethal doses of antibiotics, is responsible for most antibiotic treatment failure in MRSA infections. To understand how antibiotic tolerance is induced, *S. aureus* biofilms exposed to multiple anti-MRSA antibiotics (vancomycin, ceftaroline, delafloxacin, and linezolid) were examined using both quantitative proteomics and transposon sequencing. These screens indicated that arginine metabolism is involved in antibiotic tolerance within a biofilm and led to the hypothesis that depletion of arginine within *S. aureus* communities can induce antibiotic tolerance. Consistent with this hypothesis, inactivation of *argH,* the final gene in the arginine synthesis pathway, induces antibiotic tolerance under conditions in which the parental strain is susceptible to antibiotics. Arginine restriction was found to induce antibiotic tolerance via inhibition of protein synthesis. Finally, although *S. aureus* fitness in a mouse skin infection model is decreased in an *argH* mutant, its ability to survive *in vivo* during antibiotic treatment with vancomycin is enhanced, highlighting the relationship between arginine metabolism and antibiotic tolerance during *S. aureus* infection. Uncovering this link between arginine metabolism and antibiotic tolerance has the potential to open new therapeutic avenues targeting previously recalcitrant *S. aureus* infections.

**Significance Statement:** Methicillin-Resistant *Staphylococcus aureus* (MRSA) is a leading bacterial cause of morbidity and mortality worldwide. Despite the availability of numerous antibiotics with *in vitro* efficacy against MRSA, there are still high rates of antibiotic treatment failure in *S. aureus* infections, suggesting antibiotic tolerance is common during human infections. Here, we report a direct connection between the metabolism of arginine, an essential amino acid in *S. aureus*, and tolerance to multiple classes of antibiotics. This represents a key pathway towards broad antibiotic tolerance in *S. aureus* and therefore an attractive target to help repotentiate current antibiotics and potentially reduce treatment failure.

## Introduction

*Staphylococcus aureus* is one of the leading bacterial causes of mortality in the world (1), with mortality rates in excess of 20% for certain types of infections (2–9). These high mortality rates are due, in part, to high rates of antibiotic treatment failure that occur during the treatment of *S. aureus* infections. Anti-staphylococcal penicillin or first-generation cephalosporins are first-line treatment options for *S. aureus* infections. Although Methicillin-resistant *S. aureus* (MRSA) strains with resistance to these first-line agents are relatively common, the rates of resistance to anti-MRSA antibiotics remains very low (10, 11). In this context, the high rates of antibiotic treatment failure are surprising and suggest a mechanism besides antibiotic resistance. Multiple studies have investigated potential causes of antibiotic treatment failure in *S. aureus* and have identified a variety of contributory factors including the formation of small colony variants (SCVs), persister cells, and biofilms (12–19).

Growth as a biofilm, a dense community where adherent microbes secrete a complex extracellular matrix, induces extremely high levels of antibiotic tolerance. Antibiotic tolerance is the ability of a bacterial population to withstand an otherwise lethal antibiotic dose due to phenotypic changes without any evidence of a change in the minimum inhibitory concentration (MIC) against that antibiotic (20). Bacteria growing in a biofilm community differ from planktonic bacteria in their metabolism and growth, and they are able to tolerate 100 to 1000 times the concentration of antibiotics that would eliminate planktonic bacteria (21). Biofilm formation has been implicated in many different types of *S. aureus* infections including osteomyelitis, prosthetic joint infections, endocarditis, and chronic wound infections (22). In these infections, biofilm growth contributes to the high morbidity and recalcitrance to antibiotic treatment.

Despite much investigation and speculation about the potential causes of antibiotic tolerance in biofilm-mediated infections, the mechanisms by which this occurs in *S. aureus* are still poorly understood. In this work, an *in vitro* model of *S. aureus* biofilms grown at a solid-air interface was employed to investigate antibiotic tolerance during biofilm growth. Mechanisms of antibiotic tolerance in *S. aureus* were identified using two broad, unbiased, complementary screening approaches: semi-quantitative proteomics, and transposon sequencing-based screening. These screens identified a novel role for arginine metabolism as a key potentiator of antibiotic tolerance in *S. aureus*. By restricting the synthesis of arginine, *S. aureus* can induce antibiotic tolerance by inhibition of protein synthesis. Furthermore, inhibiting the ability of *S. aureus* to produce arginine from citrulline during antibiotic treatment enhances bacterial fitness during antibiotic treatment in a mouse model of skin and soft tissue infection (SSTI). Together, these studies demonstrate that restricting arginine synthesis, and in turn limiting arginine availability, can contribute to antibiotic treatment failure in *S. aureus*.

## Results

### Antibiotic exposure results in differences in protein abundance and relative fitness of transposon mutants in arginine metabolism pathways in *S. aureus*

To screen for proteins that are involved in antibiotic tolerance in *S. aureus* biofilms, untargeted, label-free, quantitative (LFQ) proteomics using liquid chromatography tandem mass spectrometry (LC-MS/MS) was performed. For LFQ proteomic analysis, *S. aureus* JE2, a derivative of the MRSA USA300 LAC strain, was grown in a colony filter biofilm model. This model allows for the establishment of a mature biofilm at a solid-air interface which can be easily transferred to different growth conditions while keeping the biofilm structure intact (23). Utilizing this model, *S. aureus* biofilms grown on polycarbonate filter discs on tryptic soy agar (TSA) plates could be transferred as intact biofilms to fresh media every 24 hours (Figure S1A). After 48 hours of growth in antibiotic-free conditions, mature biofilms were transferred to TSA plates containing antibiotics for an additional 48 hours. To identify pathways that are involved in tolerance to multiple antibiotics, four different classes of antibiotics were used: vancomycin, a cell wall targeting glycopeptide and the most commonly used first line antibiotic for the treatment of MRSA bacteremia worldwide (24); ceftaroline, a cell wall targeting beta-lactam antibiotic with activity against MRSA; linezolid, an oxazolidinone that inhibits protein synthesis and has activity against MRSA; and delafloxacin, a fourth generation fluoroquinolone with activity against MRSA. Bacterial killing resulting from antibiotic treatment of these biofilms after 24 and 48 hours is shown in Figure S1. After 48 hours of exposure to antibiotics or a no-antibiotic control, total protein was extracted from the colony biofilms and identified using LC-MS/MS. Based on this analysis, there were a total of 142 proteins with significant differences in their abundance when treated with one of the antibiotics tested (Table S1).

As a complementary approach to performing LFQ proteomics, a transposon library was constructed in the JE2 strain using a Himar1-based transposon approach as previously described (25). This resulted in the creation of a high quality, high-density transposon library with greater than 150,000 independent transposon insertions representing coverage of nearly 55% of all TA sites and at least one TA site in 93.3% of annotated open reading frames in the USA300_FPR3757 genome (2619 out of 2807) (Figure S2). Analysis of the library using the TRANSIT software package (26) revealed 369 essential genes in the *S. aureus* genome with another 227 genes whose essentiality was uncertain, in line with estimates from other studies in *S. aureus* (27, 28). To screen for genes impacting survival in the presence of antibiotics, the transposon library was grown using the colony filter biofilm model and exposed to antibiotics for 48 hours, as above. Following antibiotic treatment, a 4-hour outgrowth in tryptic soy broth (TSB) as a planktonic culture was performed to enrich the population of viable bacteria. Following the outgrowth, DNA was extracted, and transposon sequencing was performed. Based on analysis of the sequencing results using TRANSIT, 157 genes were either essential or detrimental to survival in at least one of the antibiotic conditions tested (Table S2). In addition to identifying genes important for survival in the presence of antibiotics, this experiment also identified genes that significantly impacted fitness during biofilm growth (Table S3).

Analysis of the datasets resulting from the LFQ proteomics and TnSeq experiments revealed that very few protein-gene pairs were identified by both techniques. However, transposon insertions disrupting either of two genes encoded in an operon, *argG* and *argH,* were found to be beneficial for survival in the presence of multiple antibiotics and the corresponding encoded proteins were decreased in abundance in response to treatment with all the antibiotics tested (Figure 1). Together, ArgG and ArgH are responsible for the synthesis of L-arginine from L-citrulline (Figure 1A). Evaluation of several other enzymes involved in arginine metabolism, ArgD, ArgC, ArgJ, ArgB, and RocD, did not show any significant differences in the proteomic or TnSeq datasets. However, the enzymes responsible for degrading arginine via the arginine deiminase pathway, ArcA, ArcB, and ArcC, showed increased abundance during exposure to 3 out of the 4 antibiotics tested (Figure 1B). Transposon insertions in *arcA, arcB,* and *arcC* did not lead to any significant fitness differences (Figure 1C). Together, these results suggest a coordinated metabolic response leading to increased arginine degradation and decreased arginine synthesis occurs in response to antibiotics during biofilm growth.

**Figure 1.**
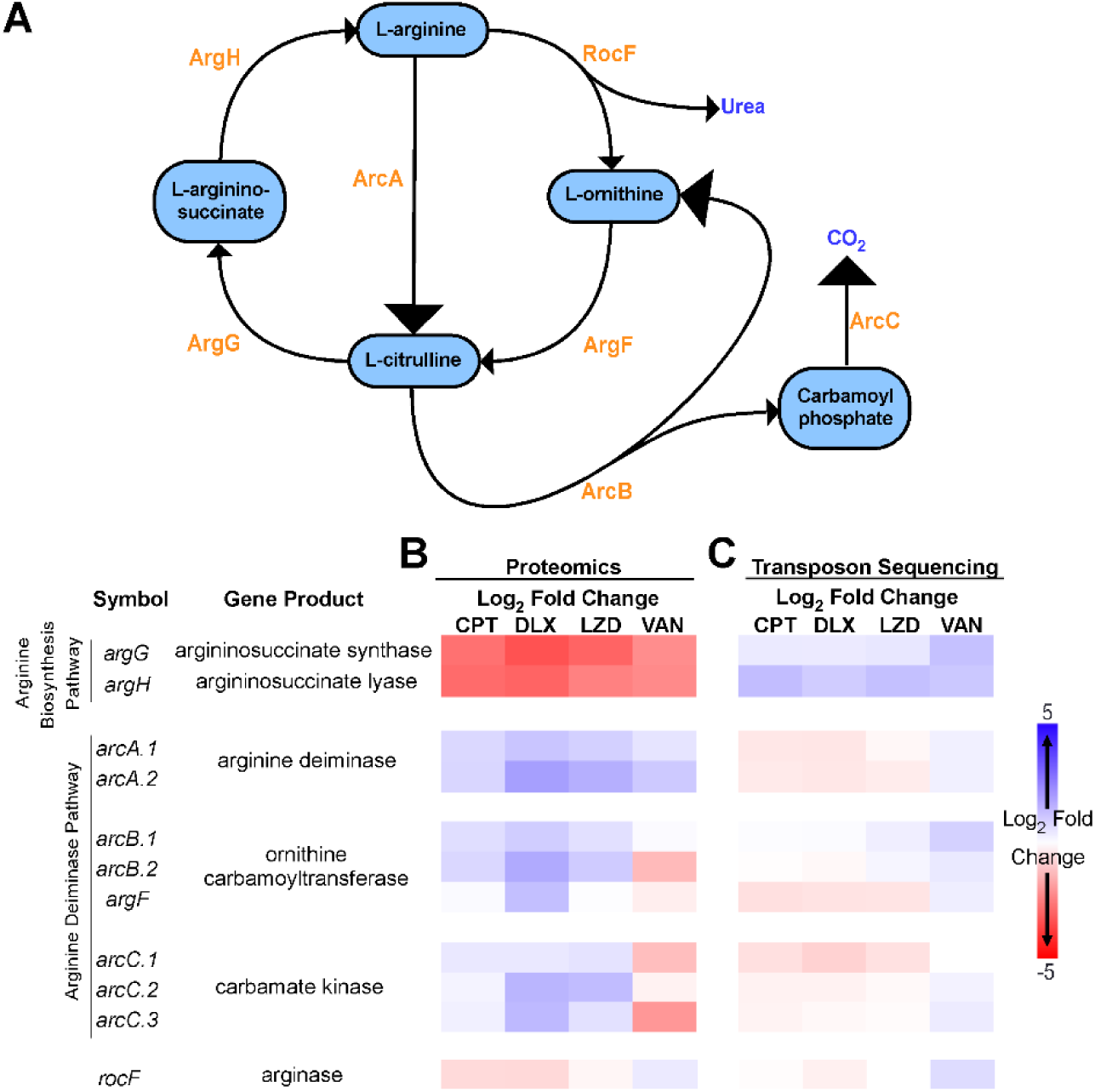
Arginine metabolism in *S. aureus* biofilms during antibiotic exposure. (**A**) Diagram showing the flux of arginine in *S. aureus,* which involves the urea cycle and the arginine deiminase pathway. The responsible enzymes for each step are shown in orange. (**B**) Mature (48 hr) *S. aureus* (strain JE2) colony biofilms grown on polycarbonate filters placed on TSA plates were transferred to fresh TSA plates either with vehicle or with one of the indicated antibiotics added. Total protein was isolated after 48 hours of exposure to the antibiotic containing media, and relative protein abundance was determined by label free quantitative LC-MS/MS proteomics. The heat map shows the *z*-scores of the log_2_ fold difference in the abundance of the indicated proteins involved in arginine metabolism after antibiotic exposure compared to the no antibiotic control. (**C**) A transposon mutant library was constructed in the JE2 background and used to grow colony biofilms. The colony biofilms were transferred to TSA plates with or without antibiotics for 48 hours after which point the biofilms were harvested and after a short outgrowth in TSB total genomic DNA was extracted for transposon sequencing. Heat map shows the *z*-scores of the log_2_ fold difference in the normalized read counts for the indicated genes involved in arginine metabolism after antibiotic exposure compared to the no antibiotic control. All experiments were done in biological triplicates. VAN=vancomycin, CPT=ceftaroline, DEL=delafloxacin, LZD=linezolid.

### Arginine is required for growth and limited within a *S. aureus* biofilm

Since arginine metabolism was implicated as having a role in antibiotic tolerance in both screens, we sought to better understand the role of arginine within *S. aureus* biofilms. *S. aureus* is unique in that it contains intact copies of the genes encoding all of the enzymes necessary to synthesize arginine from glutamate or proline, but is auxotrophic for arginine during planktonic growth (29–32). Given its requirement for arginine during planktonic growth, we hypothesized that exogenous arginine was also required for growth in a biofilm. Consistent with the phenotype reported for planktonic growth in those previous studies, JE2 was unable to grow when inoculated as a biofilm on chemically defined media lacking arginine (CDM-R) (Figure 2A). Likewise, when a 48-hour old colony filter biofilm was transferred to CDM-R, it not only was unable to grow, but it had decreased survival (Figure 2B). To determine the availability of amino acids in *S. aureus* biofilms, amino acids were extracted from 48-hour old colony filter biofilms and sent to the VUMC Analytic Services Core for analysis. Amino acid analysis of biofilms grown on both TSA and CDM (containing arginine) revealed that, even when arginine is present in the growth media, the level of free arginine in the biofilm is undetectable (Figure 2C). Collectively, this suggests that exogenous arginine is essential for growth in *S. aureus*. Furthermore, its availability is likely one of the growth-limiting factors within a biofilm, since all other essential amino acids for *S. aureus* were detected in at least one of the two media conditions (Figure 2C).

**Figure 2.**
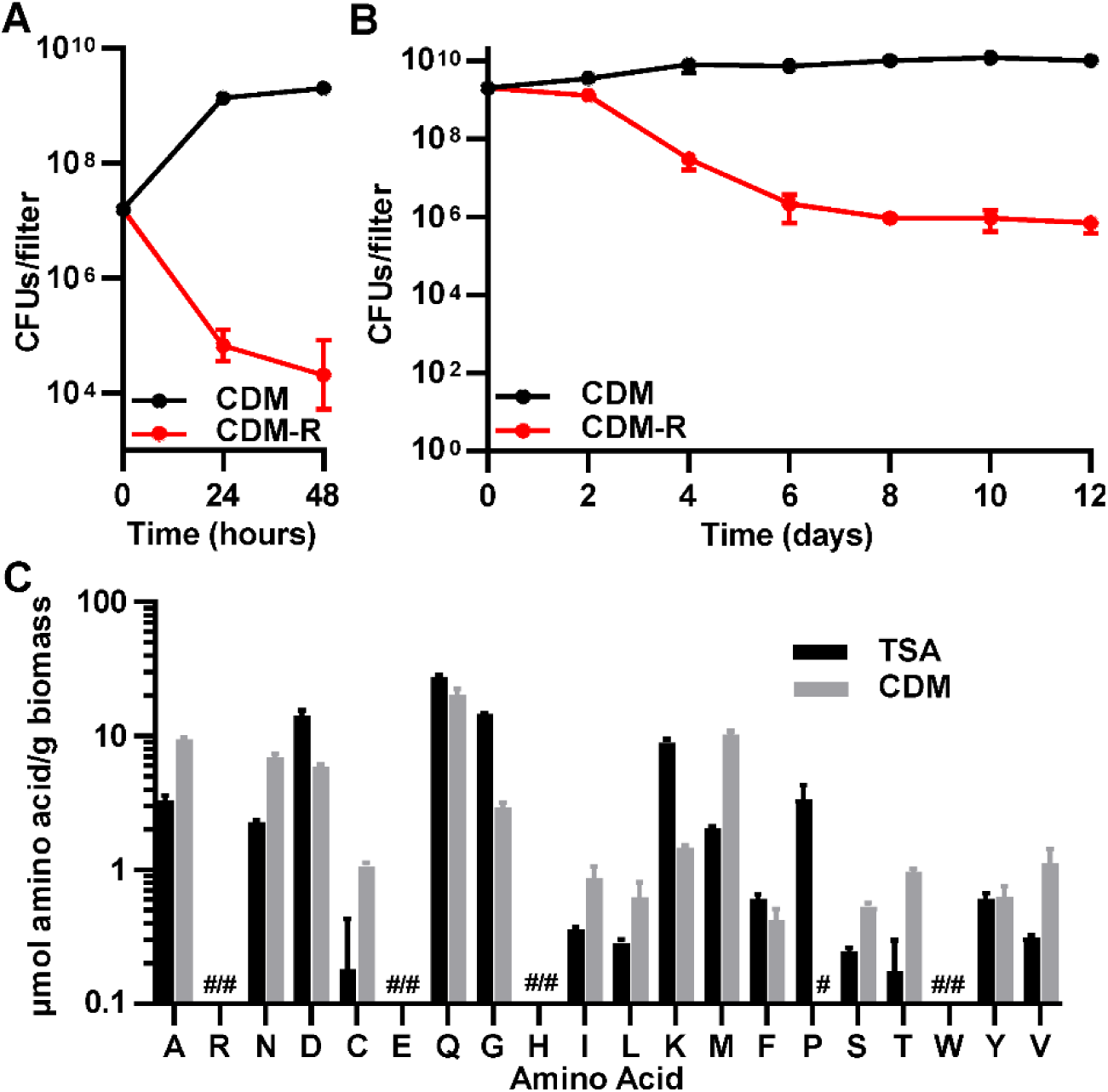
Arginine is important for growth and survival in *S. aureus* biofilms. (**A**) JE2 is unable to establish colony biofilms when inoculated on filters on chemically-defined media (CDM) in the absence of arginine, and (**B**) mature (48 hr) biofilms grown on chemically-defined media with arginine present have reduced survival when transferred to media lacking arginine (CDM-R). (**C**) Arginine levels are below the limit of detection in *S. aureus* biofilms grown on either TSA or CDM agar plates. # = below the limit of detection. Data represent technical replicates of biological triplicates.

### Restriction of arginine induces antibiotic tolerance

To understand whether arginine availability influences antibiotic tolerance, *S. aureus* was grown as colony filter biofilms on CDM for 48 hours and then the intact biofilms were transferred to either CDM or CDM-R with or without antibiotics added (Figure 3A, C, E, and G). When arginine was present in the media, all four of the antibiotics led to least a 1-log reduction in CFUs by 72 hours, when compared to the starting CFU. This was a significant reduction when compared to the untreated biofilms for all four of the antibiotics. When biofilms were transferred to media without arginine, there was a decrease in CFUs even in the absence of antibiotics. However, the addition of antibiotics to the media without arginine did not cause any further decrease in the number of CFUs when compared to the untreated biofilms, suggesting there was no effect from antibiotic treatment under arginine-restricted conditions. The only exception to this was delafloxacin, where only after 72 hours of antibiotic exposure in the absence of arginine was there a significant decrease in CFUs compared to the untreated biofilms (Figure 3E). However, this reduction in CFUs was still less than the reduction seen in biofilms treated with delafloxacin in the presence of arginine.

**Figure 3.**
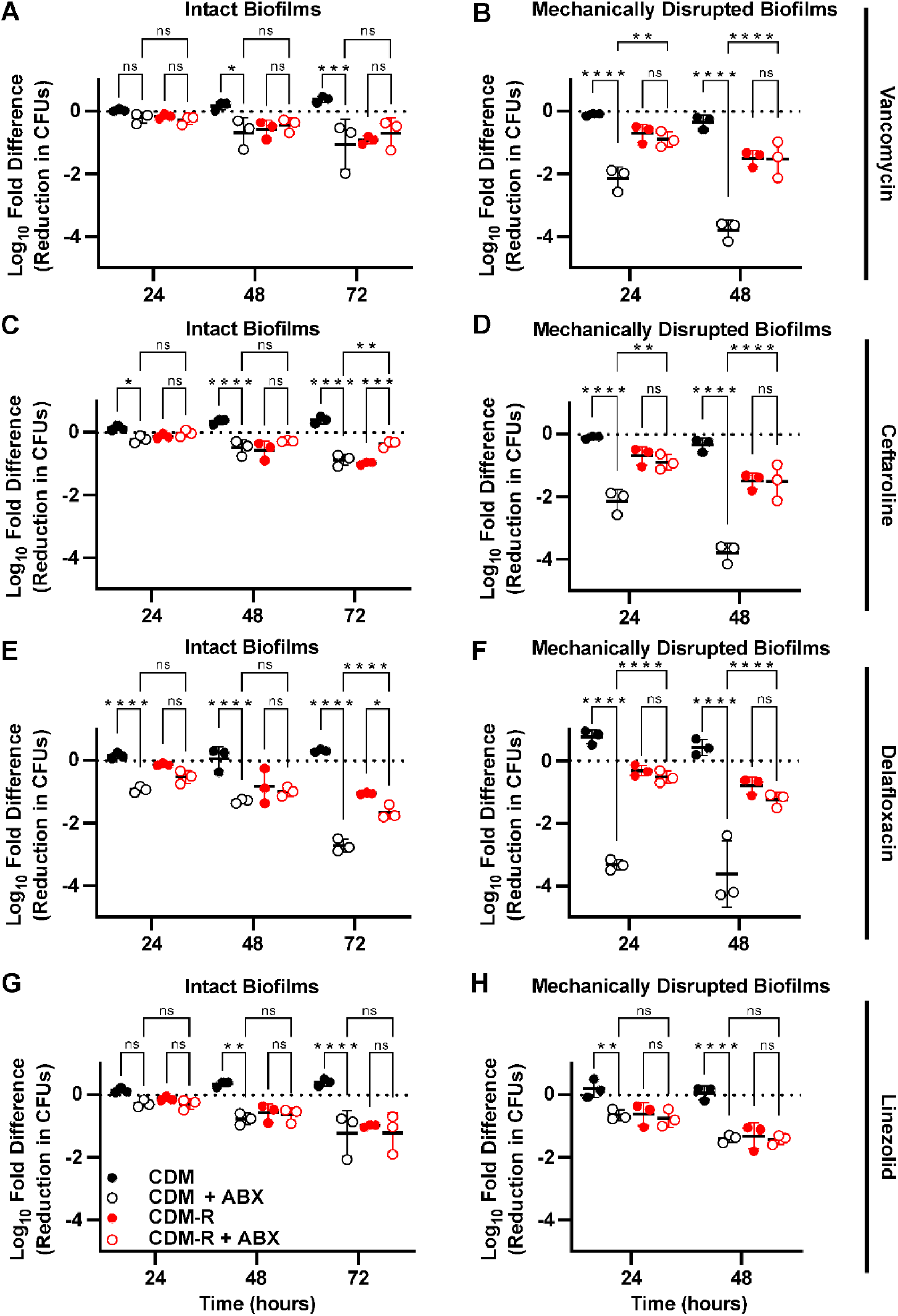
Arginine deprivation induces tolerance to multiple classes of antibiotics in *S. aureus* biofilms. Colony biofilms grown for 48 hours on polycarbonate filters on CDM agar plates were either transferred to fresh CDM or CDM-R plates with or without antibiotics (**A, C, E, G**), or were homogenized and transferred to liquid CDM or CDM-R media with or without antibiotics added (**B, D, F, H**). The following concentration of antibiotics were used: 400 µg/ml vancomycin (**A,B**), 20 µg/ml ceftaroline (**C,D**), 9 µg/ml delafloxacin (**E,F**), and 20 µg/ml linezolid (**G,H**). Data represent technical replicates of biological triplicates. 2-way ANOVA with Tukey multiple comparisons test; *=*p*<0.05, **=*p*<0.005, ***=*p*<0.0005, ****=*p*<0.0001, ns=not significant, ABX=antibiotics.

To determine if the effect of arginine on antibiotic tolerance was specific to growth in a biofilm, *S. aureus* was grown planktonically in shaking liquid culture, harvested during its logarithmic growth phase, washed, and transferred to either CDM or CDM-R with antibiotics (Figure S3). In planktonic culture, the absence of arginine only induced substantial antibiotic tolerance against ceftaroline (Figure S3B). By contrast, vancomycin, delafloxacin, and linezolid all showed greater than 2-log reductions in the number of CFUs after 48 hours of antibiotic exposure, even in the absence of arginine. Despite this, the presence of arginine in planktonic cultures did lead to a significant increase in killing by vancomycin. Differences in susceptibility to delafloxacin, however, varied over time with significantly more killing in the absence of arginine by 48 hours.

Since high concentrations of arginine weaken the integrity of biofilms in some bacterial species (33), we hypothesized that the observed effect of arginine might be due to changes in the extracellular matrix or increased antibiotic penetration within the biofilm (33). To test this hypothesis, 48-hour colony filter biofilms were homogenized, washed with PBS, and resuspended in either CDM or CDM-R broth. The homogenized biofilms were then exposed to antibiotics. Mechanically disrupted biofilms exhibited greater susceptibility to antibiotics overall when compared to intact biofilms. However, in the disrupted biofilms there was an even more pronounced difference in the amount of antibiotic killing based on the presence or absence of arginine (Figure 3B, D, F, and H). For vancomycin, ceftaroline, and delafloxacin there as significantly more antibiotic tolerance when arginine was absent. After 48 hours of antibiotic exposure, for these three antibiotics there was a greater than 100-fold difference in the number of CFUs between cultures with and without arginine. This increase in antibiotic tolerance in the absence of arginine was not restricted to JE2, as a similar increase in tolerance to vancomycin was seen with both the laboratory MSSA strain Newman and a clinical MRSA isolate (Figure S4). In homogenized JE2 biofilms, however, there was no difference in bacterial killing between the cultures with and without arginine when they were treated with linezolid, with both conditions having less than a single log reduction in CFUs. These experiments suggest an effect of arginine on antibiotic susceptibility that is dependent on the metabolism of *S. aureus* during biofilm growth, but independent of the biofilm structure.

### Restriction of arginine increases antibiotic tolerance through the inhibition of protein synthesis

The finding that, as opposed to the three other antibiotics, *S. aureus* biofilms display high levels of tolerance to linezolid regardless of arginine concentrations was intriguing. This led us to hypothesize that a pathway affected by both arginine depletion and linezolid might be responsible for the induction of antibiotic tolerance. Since linezolid is a protein synthesis inhibitor, inhibition of protein synthesis was hypothesized to be a shared pathway to induce tolerance. Although linezolid is classified as a bacteriostatic antibiotic, the concentration used in this study was sufficient to cause over a 2-log reduction in CFUs in planktonic cultures in either TSB or CDM (Figure S1 and S3). To confirm that restriction of arginine leads to inhibition of protein synthesis, nascent protein labeling was performed using click chemistry. Biofilms grown for 48 hours on CDM agar were homogenized and transferred to CDM broth lacking arginine in which L-methionine had been replaced with the methionine analog L-homopropargylglycine (L-HPG). After 4 hours bacteria were harvested, and nascent proteins were labeled to allow for visualization and quantification via western blot (Figure 4A). Normalization of the integrated density of the fluorescence signal for each sample by the total protein (Figure S5) confirmed that there was significant inhibition of protein synthesis in the absence of arginine (Figure 4B). Furthermore, the addition of citrulline reversed this inhibition of protein synthesis, presumably due to the conversion of citrulline to arginine via the ArgGH enzymes.

**Figure 4.**
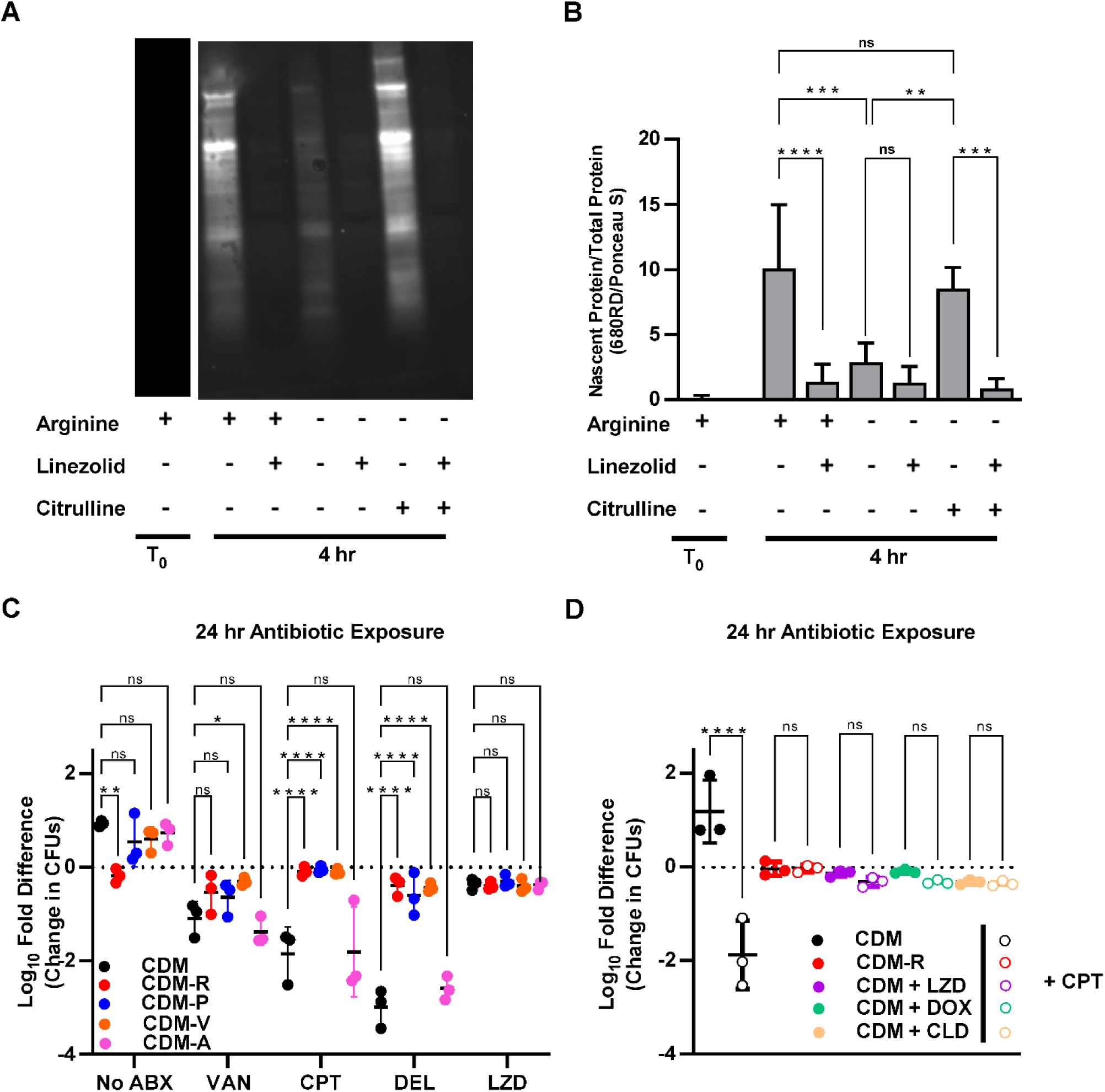
Antibiotic tolerance is mediated by amino acid starvation leading to protein synthesis arrest. (**A**) Homogenized biofilm cultures were transferred to CDM broth or CDM broth lacking individual amino acids and antibiotics were added as indicated. The reduction in CFUs compared to the starting inoculum was calculated after 24 hours of antibiotic exposure. (**B**) Homogenized biofilm cultures were transferred to fresh CDM or CDM-R and protein synthesis inhibitors were added as indicated along with either ceftaroline or a vehicle control. The reduction in CFUs compared to the starting inoculum was calculated after 24 hours of antibiotic exposure. (**C**) Representative western blot showing the labelling of nascent protein by click chemistry 4 hours after the transfer of homogenized biofilm cultures to the indicated growth conditions. (**D**) A ratio of nascent protein to total protein was calculated using integrated density values obtained by analyzing the western blots in imageJ. Data represent technical replicates of biological triplicates. 2-way ANOVA with Tukey multiple comparisons test; *\*=p<*0.05, **=*p*<0.005, ***=*p*<0.0005, ****=*p*<0.0001, ns=not significant, ABX=antibiotics, VAN=vancomycin, CPT=ceftaroline, DEL=delafloxacin, LZD=linezolid, DOX=doxycycline, CLD=clindamycin.

To validate that inhibition of protein synthesis is the mechanism by which arginine depletion leads to antibiotic tolerance, two other experiments were performed. The first experiment tested whether the depletion of other amino acids for which *S. aureus* is known to display auxotrophy also induces antibiotic tolerance. Similar to what was observed in media without arginine present, the removal of either valine or proline, two essential amino acids that *S. aureus* is unable to normally synthesize (29), increased the tolerance of *S. aureus* biofilms to vancomycin, ceftaroline, and delafloxacin (Figure 4C). By contrast, removal of the non-essential amino acid alanine had no impact on antibiotic tolerance. Notably, proline and valine could be detected in biofilms grown in TSA, albeit at low levels (Figure 2C), suggesting that these amino acids are not restricted to the degree that arginine is within *S. aureus* biofilms.

As a secondary experiment to validate that protein synthesis inhibition leads to antibiotic tolerance, the ability of multiple protein synthesis inhibitors to induce antibiotic tolerance was tested. Biofilms grown for 48 hours on CDM agar were homogenized and transferred to liquid media either lacking a protein synthesis inhibitor or containing one of three protein synthesis inhibitors (linezolid, doxycycline, or clindamycin). The addition of any one of these antibiotics that inhibit protein synthesis resulted in increased tolerance to ceftaroline, similar to what was seen in media lacking arginine (Figure 4D). Together, these experiments suggest inhibition of protein synthesis through multiple pathways, including the depletion of arginine, induces antibiotic tolerance in *S. aureus* biofilms.

### ArgGH-mediated conversion of citrulline to arginine can reverse arginine-deprivation mediated antibiotic tolerance and contributes to antibiotic susceptibility *in vivo*

Since the addition of citrulline to CDM broth lacking arginine could restore protein synthesis, it was next hypothesized that the addition of citrulline could reverse the antibiotic tolerance observed when arginine was depleted. When grown in planktonic culture, citrulline rescued the growth of JE2 in media lacking arginine but could not do so for *argH*::Tn, a strain of JE2 in which the *argH* gene was disrupted by a transposon insertion (Figure S6). As expected, the addition of citrulline reversed the antibiotic tolerance seen when arginine was absent from the media (Figure 5A). This effect was most likely due to the conversion of citrulline to arginine as citrulline did not restore antibiotic susceptibility when the experiment was repeated using *argH*::Tn (Figure 5B). Furthermore, when the *argGH* operon was reintroduced into the chromosome of the *argH*::Tn strain under a constitutively active promoter, *S. aureus* was once again able to utilize citrulline for growth (Figure S6) and also showed increased antibiotic susceptibility in the presence of citrulline (Figure 5C).

**Figure 5.**
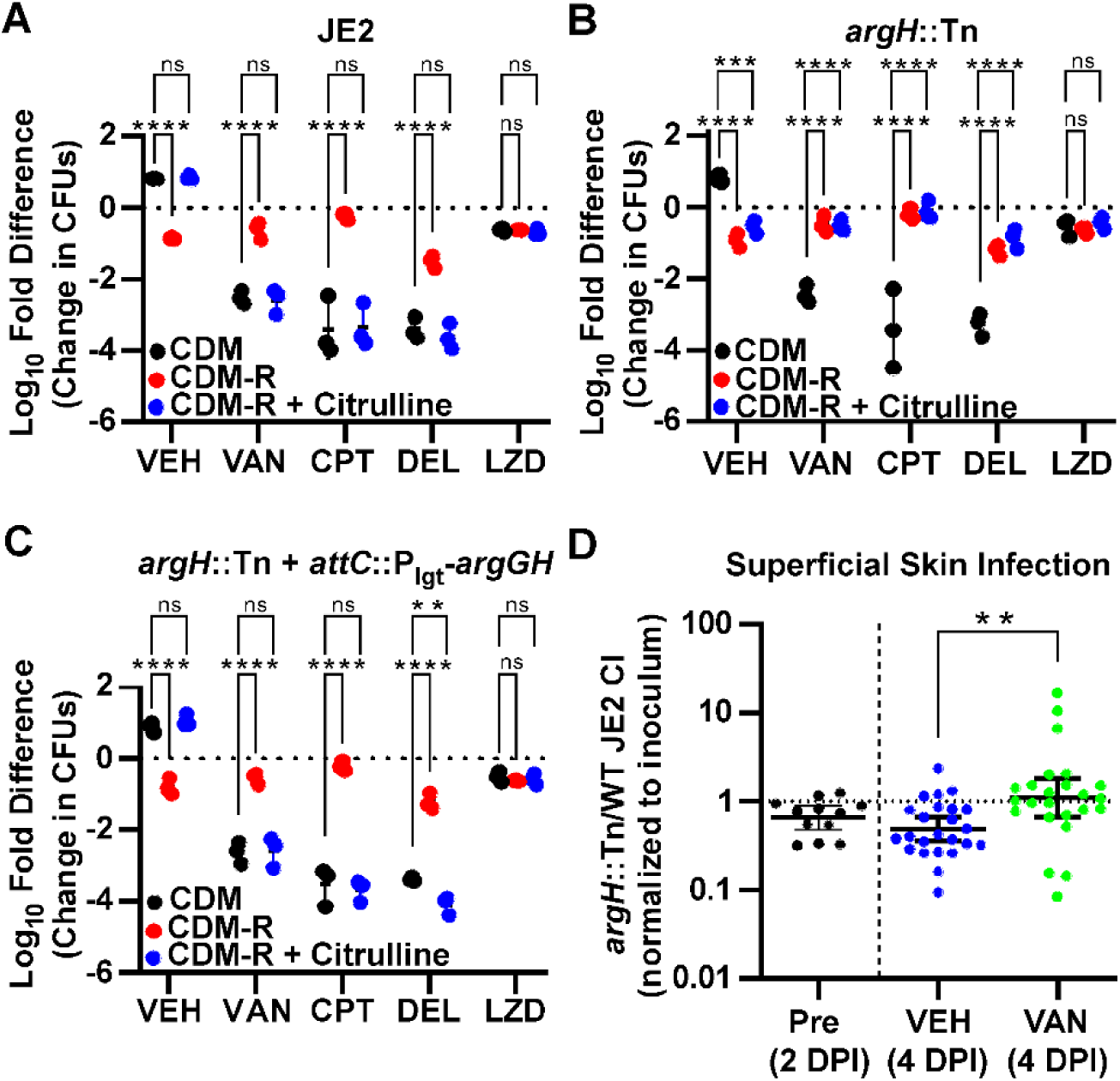
Addition of citrulline restores antibiotic susceptibility in an ArgH dependent manner. (**A**) Addition of citrulline to CDM-R restores the antibiotic susceptibility of homogenized colony biofilms to CDM levels. (**B**) Antibiotic susceptibility is not restored by the addition of citrulline in an *argH*::Tn mutant. (**C**) Complementation with the *argGH* operon in the *argH*::Tn mutant restores antibiotic susceptibility to wild-type levels in the presence of citrulline. Data represent technical replicates of biological triplicates. (**D**) Competitive index (CI) for *argH*::Tn/JE2 competition experiment in a murine superficial skin infection model. Data shown are the ratio of *argH*::Tn to JE2 CFUs, normalized to the ratio of the starting inoculum, at 2 days post infection (DPI) prior to any treatment along with the ratio after 48 hours of vancomycin or vehicle control treatment (4 DPI). Data represent the combined results from a total of 60 mice (30 females and 30 males). (**A-C**) 2-way ANOVA with Tukey multiple comparisons test, (**D**) Mann-Whitney test; *=*p*<0.05, **=*p*<0.005, ***=*p*<0.0005, ****=*p*<0.0001, ns=not significant, VEH= no antibiotic vehicle control, VAN=vancomycin, CPT=ceftaroline, DEL=delafloxacin, LZD=linezolid.

Chronically infected wounds have been shown to have elevated levels of citrulline, presumably due to metabolism of arginine by the host immune system (34). Since disruption of *argH* resulted in the inability of *S. aureus* to convert citrulline into arginine and a subsequent increase in antibiotic tolerance *in vitro*, it was hypothesized that ArgH might play an important role in antibiotic susceptibility during treatment of a *S. aureus* wound infection. Using a murine model of a skin and soft tissue infection (SSTI) (35), the ability of the *argH*::Tn strain to survive antibiotic treatment was compared directly to that of the parental strain, JE2. A patch of skin was exposed on the back of mice by tape-stripping, and the exposed skin was then inoculated with a mixture of both the JE2 and *argH*::Tn strains at a 2:1 WT:mutant ratio. The SSTI was allowed to progress for 48 hours, at which point mice received either antibiotic treatment with IP injections of vancomycin or a vehicle control. After 48 hours of treatment, mice were euthanized, and individual lesions were excised to quantify the number of CFUs of each strain present. As a control, a subset of mice was harvested at 48 hours post-inoculation (prior to any antibiotic treatment) to determine relative fitness of the two strains in the absence of antibiotics. As shown in Figure S7, there were large differences in the response to a *S. aureus* skin infection between male and female mice, with female mice showing a significant reduction in the number of total CFUs even in the absence of antibiotic treatment. However, the changes within the ratio of mutant to wildtype (as measured by a competitive index), were relatively consistent across both sexes. Among all mice, in the absence of antibiotic treatment, there was a significant decrease in fitness of the *argH:*Tn mutant relative to the parental control at both 2 DPI and 4 DPI (one-sample Wilcoxon test, *p*=0.0269 and *p*=0.0008, respectively), in line with previous studies showing a virulence defect in an *argH* mutant during an infection (30). Conversely, the *argH* mutant had a higher competitive index in the vancomycin treatment group when compared to either the pretreatment or vehicle control treated groups (Figure 5D), consistent with the hypothesis that lower levels of ArgH are beneficial to *S. aureus* during antibiotic treatment. These experiments support an important role for the conversion of citrulline into arginine by ArgGH in influencing antibiotic tolerance during an infection.

## Discussion

Through the experiments detailed above, we uncovered a previously unappreciated relationship between arginine availability, arginine metabolism, protein synthesis, and antibiotic tolerance in *S. aureus*. This relationship was elucidated with the help of two broad screening approaches carried out in parallel, LFQ proteomics and TnSeq. Although, with the exception of the enzymes involved in arginine metabolism, there were very few gene/protein pairs identified as hits in both datasets, both techniques provide important insight and complementary information. Both screens independently identified different sets of genes that have been previously shown to influence antibiotic susceptibility (36–42). As an example, the proteomics approach is useful for identifying changes in protein abundance that may be missed through a transposon screen due to functional redundancy or compensatory mechanisms. This likely explains why components of the arginine deiminase pathway were identified in the proteomics screen, but not in the transposon screen. Conversely, our transposon screen can identify effects related to lower abundance proteins that cannot be accurately quantified via LFQ proteomics or proteins whose functions are controlled by post-translational regulation or other mechanisms that do not involve changes in total abundance to exert their influence. This is the case for many genes identified as contributing to antibiotic tolerance in our transposon screen (*graXRS, arlRS, mprF*, and *vraFG*, among others) that did not show significant differences or were not found in the proteomic dataset, but are already known to influence antibiotic susceptibility in *S. aureus* during planktonic growth (39–42).

Our experimental design also allowed us to identify genes that were required for biofilm growth (Table S3). However, since genes required for survival in a biofilm were selected against by the 48 hours of biofilm growth that occurred prior to antibiotic exposure, we were unable to test their contribution to antibiotic tolerance directly. It is likely that many of these genes that contribute to biofilm fitness also play a role in antibiotic tolerance and may warrant further investigation. As an example, VraSR, the <u>v</u>ancomycin-resistance-<u>a</u>ssociated two component system is known to be associated with susceptibility to vancomycin (36) and the proteomics experiments showed significant increases in levels of VraS, VraR, and the majority of the proteins known to make up the VraSR regulon (Table S1)(43). However, *vraR* and *vraS* mutants were found to be essential for biofilm growth (Table S3), and therefore not identified as having decreased fitness in the presence of vancomycin in our transposon screen. This highlights one of the benefits of our complementary screening approach. A similar explanation may explain why genes such as *ychF, ndh2, spsA, addA, purE, bfmBAB,* and *sgtB*, all of which were increased in abundance in the proteomic screen and essential for biofilm growth in our transposon screen, were not identified as playing a role in fitness during antibiotic exposure in our transposon sequencing experiment.

While an unanticipated finding from our screen, a connection between antibiotic tolerance and arginine metabolism is not entirely unprecedented. Studies in other bacterial species have found that arginine and arginine metabolism can impact antibiotic susceptibility, including during biofilm growth (44–47). Although arginine can disrupt the extracellular matrix of bacterial biofilms at very high concentrations (33), arginine’s primary effect on antibiotic tolerance in *S. aureus* is not through disruption of the biofilm. To exclude the possibility that arginine affects antibiotic susceptibility by disrupting extracellular matrix, biofilms were homogenized for the majority of the experiments in this work. In these experiments, biofilm bacteria were resuspended in shaking liquid cultures at a high concentration to preserve the high density, nutrient-limited conditions found in a biofilm. This technique also effectively removed any antibiotic tolerance that was the result of variable penetration of an antibiotic within a biofilm from our experiments. Not only was a difference between media with and without arginine preserved under these conditions, but the difference in antibiotic susceptibility was enhanced for all the antibiotics tested. The ability of linezolid and other antibiotics that inhibit protein synthesis to induce antibiotic tolerance similar to what was seen in arginine depletion (Figure 4D), supports inhibition of protein synthesis as a common mechanism. Inhibition of protein synthesis from amino acid starvation is known to induce the stringent response in *S. aureus* (48). Since activation of the stringent response can induce tolerance to antibiotics (49), this mechanism likely explains some, if not the majority, of the means by which arginine depletion ultimately leads to antibiotic tolerance (49).

There has been a growing appreciation for the role of antibiotic tolerance in *S. aureus* treatment failure which has coincided with the uncovering of multiple mechanisms by which tolerance is induced. Depletion of ATP, inhibition of the TCA cycle by reactive oxygen species (ROS), and induction of the stringent response have all been tied to antibiotic tolerance in *S. aureus* (50–53). It is not yet clear where arginine depletion fits within these other mechanisms. However, it is intriguing that arginine has direct ties to these other pathways, as not only is arginine deprivation likely to induce the stringent response through amino acid starvation but arginine can be used as an alternative source of ATP production (54). Furthermore, arginine is essential for the production of the ROS nitric oxide by host macrophages (55), suggesting that production of ROS by the host immune system may be tied directly to the depletion of arginine during an infection. While outside the scope of these studies, it would be interesting to investigate the connection between arginine depletion and tolerance specifically within the context of interaction with host immune cells.

The ability of *S. aureus* to induce tolerance by restricting its own arginine synthesis pathway may explain a long-standing paradox-namely that wildtype *S. aureus* has fully functional arginine synthesis enzymes yet does not synthesize arginine under any *in vitro* conditions that have been tested thus far (31, 32). In line with other studies, *S. aureus* was unable to grow without arginine (Figure 2). Among the enzymes in the arginine synthesis pathway, only ArgG and ArgH were found at levels above the limit of detection in our proteomic dataset. ArgD, ArgC, ArgJ, and ArgB were not found, consistent with other work showing these enzymes are under high levels of transcriptional repression (30–32, 56). *S. aureus* contains multiple enzymatic pathways that catabolize arginine, which helps create an environment that can rapidly consume any exogenous arginine, thus limiting the availability of arginine for protein synthesis. These pathways likely maintain a low baseline level of arginine in *S. aureus* biofilms, as evidenced by the inability to detect arginine within mature biofilms (Figure 2C). The isolate used in this study, JE2, is a representative USA300 strain, which has become the most common *S. aureus* strain type in the United States (57). Intriguingly, almost all USA300 strains have two full copies of the arginine deiminase pathway due to a second copy that is contained within the Arginine Catabolism Mobile Element (ACME) (58). The acquisition of the ACME has been postulated to be related to the success of USA300 strains, although a connection has not been clearly established. This second arginine deiminase pathway may be contributing to the success of USA300 through a role in antibiotic tolerance, a hypothesis furthered by the fact that the ACME arginine deiminase pathway is constitutively expressed as opposed to the native arginine deiminase pathway that is only expressed under anaerobic conditions (59). While there was some variability across the proteomic dataset in the levels of the arginine deiminase pathway enzymes based on the antibiotic tested, an increase in the abundance of both copies of ArcA, the first enzyme in the pathway, were seen across all four antibiotics tested (Figure S1).

Although *S. aureus* was unable to synthesize arginine in the presence of glutamate or proline (Figure 2), it was able to utilize exogenous citrulline to synthesize arginine presumptively via ArgG and ArgH activity (Figure S6). The ability of exogenous citrulline to increase antibiotic susceptibility was shown in this study both *in vitro* (Figure 5A, 5B, and 5C) and *in vivo* in a skin infection and treatment model as part of a competition experiment (Figure 5D). In this model, an *argH* mutant had a relative fitness defect compared to the parental JE2 strain, consistent with a prior *in vivo* study using the *argH* mutant (30). However, disruption of *argH* led to an increase in the relative fitness of the mutant during vancomycin treatment. Together, these studies support the idea that the conversion of citrulline to arginine through ArgH is an important source of arginine for *S. aureus.* Plasma arginine and citrulline levels are decreased in humans during sepsis (60, 61), suggesting a possible opportunity for a therapeutic intervention that improves antibiotic effectiveness by increasing levels of amino acids at the site of an infection. The uncovering of this previously underexplored connection between bacterial metabolism, arginine availability, and antibiotic tolerance represents an exciting new target to help overcome antibiotic treatment failure.

## Materials and Methods

Descriptions of the bacterial strains and growth conditions, colony filter biofilm assay, proteomic sampling, LC-MS/MS, bioinformatic analysis, transposon library construction, screen, and analysis, homogenized biofilm assay, amino acid quantification, labeling of nascent protein, and murine superficial skin infection and treatment model are in the SI Materials and Methods. All animal experiments were reviewed and approved by Vanderbilt University Medical Center (VUMC) Institutional Animal Care and Use Committee.

## Acknowledgments

Plasmids pBursa and pMG020 used in the construction of a transposon library were graciously provided by Dr. Anthony Richardson. We appreciate the assistance of Dale Chaput and the University of South Florida Proteomics & Mass Spectrometry Core Facility for assistance with the proteomics experiments. We appreciate the assistance of the VUMC Analytical Services Core (supported by NIH grants DK059637 and DK020593) with the amino acid quantification experiments. We appreciate the Tufts University Core Facility Genomics Core for assistance with the transposon sequencing experiments. J.A.F. received support from the VUMC Faculty Research Scholars program along with support from the National Institutes of Health (NIH) through a T32 training grant (AI007540) and an F32 postdoctoral fellowship (AI169905). V.M.R.R. was supported by the Howard Hughes Medical Institute as an HHMI Hanna Gray Fellow and by the Academic Pathways program from Vanderbilt University. V.M.R.R. holds a Postdoctoral Enrichment Program Award from the Burroughs Wellcome Fund. This work was funded by R01AI150701, R01AI138581, R01AI17829, and R01AI145992 to EPS. We appreciate members of the Skaar laboratory for providing critical review and feedback of this manuscript.

