## Supplemental Information for "Restriction of Arginine Induces Antibiotic Tolerance in *Staphylococcus aureus*"

### SI Materials and Methods

**Bacterial Strains and Growth Conditions.** JE2 is a previously described strain derived from the MRSA USA300 LAC (1). Newman is a previously described MSSA strain (2). *S. aureus* CI 5296 is a clinical *S. aureus* isolate that was obtained from the VUMC clinical microbiology laboratory with approval from the VUMC Institutional Review Board. CI 5296 was initially isolated from the blood culture of a patient with endocarditis. Antibiotic susceptibility testing initially performed as part of routine clinical microbiology laboratory testing identified CI 5296 as methicillin-resistant, with susceptibility to ceftaroline (MIC 0.5), linezolid (MIC ≤ 1), and vancomycin (MIC = 2). Strain *argH*::Tn was constructed by phage transduction of a*rgH*::*erm* from the Nebraska Transposon Mutant Library (NTML) strain NE106 into JE2 via φ85. Correct construction of the strain was confirmed by PCR as previously described for the NTML (1) and by whole genome sequencing.

*S. aureus* was routinely grown in planktonic culture in Tryptic Soy Broth (TSB; Becton, Dickinson and Company) at 37°C with shaking at 180 rpm when not explicitly grown as a biofilm. When indicated, antibiotics were added to growth media to the following concentrations- 400 µg/ml vancomycin (Fresenius Kabi), 20 µg/ml ceftaroline (Sigma Aldrich), 9 µg/ml delafloxacin (Sigma Aldrich), 20 µg/ml linezolid (Thermo Scientific Chemicals), 2.1 µg/ml doxycycline (Sigma Aldrich), and 14 µg/ml clindamycin (Thermo Scientific Chemicals). With the exception of vancomycin antibiotic concentrations used were chosen based on the peak serum concentrations for standard clinical treatment doses as published in the Sanford Guide to Antimicrobial Therapy (3). Chemically defined media (CDM) was prepared as described in Vitko *et. al.* (4) with the correction that the concentration of magnesium sulfate heptahydrate in the stock salt solution was 2.56 g/L (instead of 25.6 g/L). When indicated, CDM was prepared with the omission of individual amino acids, when indicated. CDM agar was prepared by combining the necessary stock salt solution, amino acids, and bases with 15 g/L agarose and autoclaving the media prior to the addition of the vitamin solution, trace elements, and any antibiotics as indicated.

**Colony Filter Biofilm Assay.** Colony biofilms were grown on membrane filters in a manner similar to Anderl *et. al.* (5). Overnight cultures of *S. aureus* were pelleted by centrifugation, washed with phosphate buffered saline (PBS), and diluted to an OD600 of 0.1. Polycarbonate membrane filters (13-mm diameter;

0.2 µm pore size) (Whatman) were placed on top of nutrient agar plates and were inoculated by placing 10 µl of the diluted overnight culture on the center of the filter. The agar plates containing the filters were incubated upside down at 37°C, and filters were transferred to fresh nutrient agar plates every 24 hours. Biofilms were homogenized by placing filters in 1.5 mL Navy Eppendorf Bead Lysis Kits (Next Advance, Inc) with 1 mL PBS and homogenizing them using a Bullet Blender Tissue Homogenizer (Next Advance, Inc) run for 4 successive cycles of 3 minutes at max speed at 4°C. Serial dilutions of the homogenates were plated to determine CFUs. For antibiotic susceptibility testing of intact biofilms, filters containing mature (48 hr old) biofilms were transferred to fresh nutrient agar plates containing the indicated antibiotics. Filters were transferred to fresh antibiotic containing nutrient agar plates every 24 hours until they were homogenized and CFUs were determined by plating serial dilutions. Each experiment was repeated in triplicate with technical replicates used each time. Data were analyzed using either Student T-test or 2-way ANOVA depending on the number of conditions being tested, with the appropriate corrections for multiple comparisons included.

**Planktonic antibiotic susceptibility testing.** To test the antibiotic susceptibility of planktonic JE2, overnight cultures were diluted 1:100 into prewarmed media (either TSB or CDM) and grown at 37°C, shaking at 180 rpm**.** After 2 hours of growth (mid-exponential phase), cultures were removed from the incubator, pelleted by centrifugation, washed with PBS, and diluted to an OD600 of 0.5 in fresh TSB. This culture was split into individual aliquots and antibiotics were added to the final concentrations outlined above. These cultures were returned to the 37°C shaking incubator for the remainder of the experiment. At 4, 24, and 48 hours, 1 mL aliquots of each culture were removed, pelleted by centrifugation, washed with PBS, and resuspended in 1 mL before being plated to determine CFUs by serial plating. In the case of experiments examining the role of arginine on planktonic antibiotic susceptibility, a similar procedure was employed with the exception that the overnight culture of JE2 was diluted 1:100 into CDM instead of TSB. For these experiments, after washing the mid-exponential cultures with PBS, the cells were resuspended in CDM without arginine. This culture was then split into two aliquots, and L-arginine was added to one aliquot to a final concentration of 400 µM L-arginine (the concentration of arginine in CDM) before the cultures were split further and antibiotics were added.

**Proteomic sampling.** Mature colony filter biofilms inoculated and grown on TSA as above were transferred to TSA plates either without antibiotics or with vancomycin, ceftaroline, delafloxacin, or linezolid added. The colony filter biofilms were allowed to grow for an additional 48 hours, with transfer to a fresh plate containing the same antibiotic growth conditions after 24 hours. After 48 hours of antibiotic exposure, the filters containing the biofilms were transferred to a microcentrifuge tube and frozen at -20°C until protein extraction could be performed. In each experiment, three colony biofilm filters were pooled for each sample, and the experiment was repeated in triplicate. Protein extraction was performed by washing the filters in SA lysis solution (50 mM TrisHCL pH7.5, 20 mM $MgCl_{2,}$ Roche Complete protease inhibitors) and vortexing to separate the biofilm from the filters and resuspend the bacteria. The samples were washed once with SA lysis solution, and then resuspended in SA lysis solution with 40 µg/ml of lysostaphin (AMBI Products) and incubated at 37°C for 1 hour. Following lysostaphin digestion, IGEPAL (Sigma Aldrich) was added to a final concentration of 1% and the samples were incubated on ice for 15 minutes prior to sonication. Following sonication, the cellular debris was pelleted by centrifugation, and the supernatant was transferred to a fresh microcentrifuge tube and stored at -20°C prior to LC-MS/MS analysis.

**LC-MS/MS.** To generate quantitative proteomics data, proteins were solubilized with 5% SDS, 50mM TEAB (pH 7.6), incubated at 95°C for 5 minutes, and sonicated at 20% amplitude. Protein concentrations were determined using the Pierce 660 Assay (Thermo Scientific), and equal amounts of protein were digested using S-traps (Protifi). Briefly, proteins were reduced with dithiothreitol (DTT), alkylated with iodoacetamide (IAA), acidified using phosphoric acid, and combined with s-trap loading buffer (90% MeOH, 100mM TEAB). Proteins were loaded onto s-traps, washed, and digested with Trypsin/Lys-C overnight at 37˚C. Peptides were eluted and dried with a vacuum concentrator. Peptides were resuspended in H2O/0.1% formic acid for LC-MS/MS analysis.

Peptides were separated using a 75 µm x 50 cm C18 reversed-phase-HPLC column (Thermo Scientific) on an Ultimate 3000 UHPLC (Thermo Scientific) with a 120-minute gradient (2-32% ACN with 0.1% formic acid) and analyzed on a hybrid quadrupole-Orbitrap instrument (Q Exactive Plus, Thermo Fisher Scientific). Full MS survey scans were acquired at 70,000 resolution. The top 10 most abundant ions were selected for MS/MS analysis.

**Proteome bioinformatic analysis.** Raw data files were processed in MaxQuant (www.maxquant.org) (6) and searched against the current Uniprot *S. aureus* protein sequences database. Search parameters include constant modification of cysteine by carbamidomethylation and the variable modification, methionine oxidation. Proteins are identified using the filtering criteria of 1% protein and peptide false discovery rate.

**Transposon library construction.** A transposon library was constructed in the JE2 strain using the plasmids pBursa and pMG020 as described in Grosser *et. al.* (7). The library construction process was repeated multiple times until enough transposon mutants could be pooled together to make a single library. This pooled library was grown for an additional 4 hours in TSB (without antibiotics) before being frozen in individual aliquots at a concentration of roughly $1x10^{11}$ CFUs/ml at -80°C. Spot plating of the inoculum on TSA plates containing antibiotics (either 10 µg/ml tetracycline or 10 µg/ml chloramphenicol 10 µg/ml) confirmed curing of plasmids pBursa and pMG020 in greater than 99.98% of bacterial cells. The library was confirmed to have roughly 150,000 independent transposon mutants as verified by Illumina sequencing analysis. Pooled aliquots of this high-density library were frozen at -80°C until used.

**Transposon library screen.** A single aliquot of the frozen JE2 transposon library was thawed on ice, diluted 1:100 in PBS, and 10 µL (corresponding to $\sim 1x10^{7}$ CFUs) of the diluted transposon library were inoculated on a polycarbonate filter to form a colony filter biofilm as above. These biofilms were grown on TSA for 48 hours prior to being transferred to TSA plates either without antibiotics or with vancomycin, ceftaroline, delafloxacin, or linezolid added. The biofilms were grown for an additional 48 hours, as above. After 48 hours of antibiotic exposure, the filters containing the biofilms were transferred to a microcentrifuge tube, washed with 1 mL PBS, and vortexing to separate the biofilm from the filters and resuspend the bacteria. The resulting suspension was then diluted 1:500 into fresh TSB without antibiotics and outgrown for 4 hours to enrich for viable bacteria. In addition to collecting samples after 48 hours of antibiotic exposure, samples were also collected prior to transfer to antibiotic containing media, to serve as a $T_0$ comparison. The resulting culture was centrifuged, and the bacterial pellet was stored at -80°C until DNA extraction could be performed. In each experiment, three colony biofilm filters were pooled for each sample, and the experiment was repeated in triplicate.

**Transposon sequencing analysis.** The frozen bacteria pellets from the transposon library screen were thawed on ice and genomic DNA was extracted using the DNeasy Blood and Tissue Kit (Qiagen). Genomic DNA was sheared to 350 bp by sonication using a Covaris LE220 and prepared by sequencing using the homopolymer tail-mediated ligation PCR (HTML-PCR) as outlined in van Opijnen *et. al.* (8) with the modification of the use of the KAPA HiFi HotStart DNA Polymerase (Roche) for the PCR cycles and the use of the transposon-specific primers olj510 and olj511 as previously outlined (7). Replicates were all individually barcoded and then multiplexed for sequencing. Sequencing was performed using a custom sequencing primer (olj512) on the HiSeq 2500 (Illumina) by the Tufts University Genomics Core Facility. Sequencing data was analyzed using the TRANSIT software package for TnSeq analysis (9). Reads were processed and mapped to the *S. aureus* FPR3757 genome using the TnSeq Pre-Processor (TPP) tool. Gene essentiality was determining using the Gumbel method in TRANSIT. $\text{Log}_2$ fold changes between antibiotic exposed biofilms and the no antibiotic controls were calculated using the Resampling method in TRANSIT with beta-geometric normalization and correction for multiple comparisons using the Benjamini-Hochberg procedure.

**Homogenized Biofilm Assay.** For assays involving mechanical disruption of biofilms, colony filter biofilms were grown for 48 hours as described above. At that time, biofilms were transferred to 1.5 mL Red Eppendorf Bead Lysis Kits (Next Advance, Inc) with 1 mL PBS and homogenizing using a Bullet Blender Tissue Homogenizer (Next Advance, Inc) run for 4 successive cycles at max speed at 4°C. The biofilm homogenate was centrifuged to pellet the bacteria, the supernatant was discarded, and the bacteria was resuspended and diluted in CDM lacking arginine and other amino acids as appropriate for a given experiment at a ratio of 2.5 mL media per filter biofilm. The homogenized biofilm culture was then split into equal aliquots to which either a vehicle control or the missing amino acids were added to the appropriate concentration. The samples were then aliquoted into a 96 well plate and antibiotics were added individual wells, as appropriate for a given experiment. The plates were incubated at 37°C with shaking at 180 rpm for up to 48 hours. In addition to plating the homogenized cultures at the time of transfer to the 96 well plate, aliquots were also removed at 24 hours and 48 hours, and serial dilutions were plated to determine CFUs remaining. Each experiment was repeated in triplicate with technical replicates used each time. Data were analyzed using either Student T-test or 2-way ANOVA depending on the number of conditions being tested, with the appropriate corrections for multiple comparisons included.

**Amino Acid Quantification.** Colony filter biofilms grown for 48 hours, as above, were weighed and stored at -80°C. Amino acid extraction was performed by washing the filters in SA lysis solution and vortexing to separate the biofilm from the filters and resuspend the bacteria. The samples were washed once with SA lysis solution, and then resuspended in SA lysis solution with 40 µg/ml of lysostaphin (AMBI Products) and incubated at 37°C for 1 hour. Following lysostaphin digestion, 5-sulfosalicylic acid (Fisher Chemical) was added to a concentration of 20%, and the samples were sonicated. Following sonication, the cellular debris was pelleted by centrifugation, and the supernatant was transferred to a fresh microcentrifuge tube and stored at -20°C prior to amino acid analysis. Amino acid concentrations were determined by HPLC by the VUMC Hormone Assay and Analytical Services Core using a dedicated Biochrom 30 amino acid analyzer.

**Labeling of Nascent Protein.** Mature colony filter biofilms were homogenized as described above but were diluted into CDM lacking both arginine and methionine. The methionine analog L-homopropargylglycine (Thermo Fisher Scientific) was added to a final concentration of 470 µM and the culture was split into three equal aliquots to which either a vehicle control, L-arginine, or L-citrulline was added to a final concentration of 400 µM. These aliquots were split further, and a vehicle control or linezolid was added to a concentration of 20 µg/ml. The resulting cultures were incubated at 37°C with shaking at 180 rpm. At 1 and 4 hours following the split of the cultures, samples were collected from each culture. Cells were pelleted by centrifugation and stored at -80°C for future use. The cell pellets were thawed on ice, washed once with SA lysis solution, and then resuspended in SA lysis solution with 40 µg/ml of lysostaphin (AMBI Products) and incubated at 37°C for 15 minutes. Following lysostaphin digestion, SDS (Fisher Chemical) was added to a final concentration of 1% and the samples were incubated on ice for 15 minutes prior to sonication. Following sonication, the cellular debris was pelleted by centrifugation, and the supernatant was transferred to a fresh microcentrifuge tube and stored at -20°C. Protein concentration in each sample was determined using a Pierce BCA Protein Assay Kit (Thermo Scientific) with SA lysis buffer containing lysostaphin used to normalize the protein concentrations. Equal concentrations of protein for each sample were used for click chemistry labeling using the Click-iT Protein Reaction Buffer Kit (Thermo

Fisher Scientific) and Biotin Azide (Thermo Fisher Scientific) according to the manufacturer's instructions. The resulting samples were resolubilized in Laemmli Sample Buffer and run on an SDS-PAGE gel prior to transfer to a nitrocellulose membrane. Total protein was stained using Ponceau S stain (Sigma-Aldrich) according to the manufacturer's instructions and then subsequently destained. The membrane was then blocked using Intercept Blocking Buffer (Li-Cor) and stained using IRDye 680RD Streptavidin (Li-Cor). Images of total protein staining with Ponceau S and nascent protein staining with IRDye 680RD Streptavidin were both captured using a ChemiDoc Imaging System (Bio-Rad) with the manufacture's preprogramed settings for Ponceau S and IRDye 680RD, respectively. The integrated density of each sample was determined using imageJ analysis.

**Murine Superficial Skin Infection and Treatment Model.** A well-established murine model of a skin and soft tissue infection was employed as previously described (10). Briefly, 6-8 week old C57BL/6 mice (Jackson Laboratories) were anesthetized using isoflurane. Tensoplast® adhesive bandages were used to remove fur from a roughly 2 cm x 2 cm patch on the back of a mouse. Overnight cultures of JE2 and *argH*::Tn were separately subcultured into TSB and grown to mid log phase, at which point they were washed with PBS and combined in an OD600-matched ratio of 2:1 (JE2 to *argH*::Tn). A 5 µL droplet of this bacterial suspension was spread on the exposed skin of the anesthetized mice and allowed to dry before the mice recovered from anesthesia. At 48 hours post infection, a subset of mice were humanely euthanized, and their lesions were excised to enumerate the number of CFUs present. Mice were then treated for 48 hours with twice daily intraperitoneal infections of either a 30 mg/kg of vancomycin (administered as a 3 mg/ml solution of vancomycin in PBS) or an equivalent volume of PBS alone (vehicle control). After 48 hours of treatment, the remaining mice were humanely euthanized, and their lesions were excised for CFU enumeration. Each lesion was placed in a 1.5 mL Navy Eppendorf Bead Lysis Kits (Next Advance, Inc) with 1 mL PBS and homogenizing them using a Bullet Blender Tissue Homogenizer (Next Advance, Inc) run for 3 successive cycles of 5 minutes at max speed at 4°C. Serial dilutions of the homogenates were plated in triplicate to determine CFUs. Samples were plated on TSA plates containing Erm 10 µg/ml to determine the number of CFUs of *argH*::Tn present and were plated on TSA plates containing Ciprofloxacin 2 µg/ml to determine the total number of CFUs of *S. aureus* present. This concentration of ciprofloxacin was used as it was separately determined to inhibit the growth of other skin flora without affecting the growth of JE2 or *argH*::Tn. The number of CFUs of JE2 present in a sample was calculated by subtracting the number of CFUs of *argH*::Tn from the total number of CFUs present. Samples were also plated on TSA without any antibiotics as a control to examine the presence of normal skin flora. These studies were approved by the VUMC Institutional Animal Care and Use Committee.

**Figure S1**

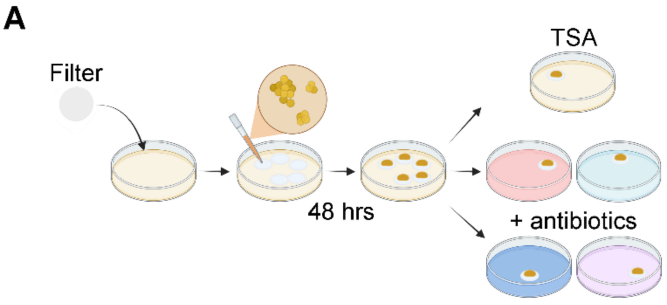


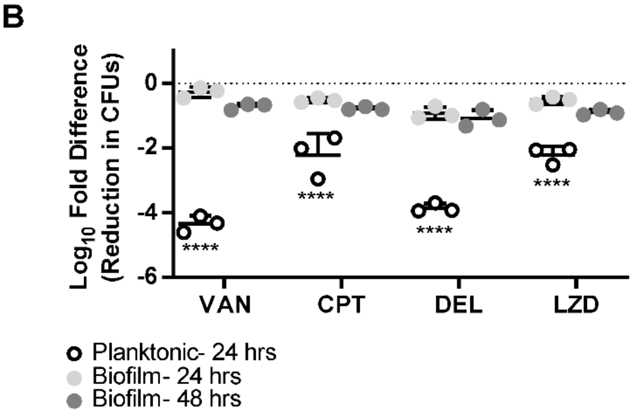


**Figure S1. Colony filter biofilms exhibit increased levels of antibiotic tolerance. (A)** *S. aureus* strain JE2 was grown for 48 hours on polycarbonate filters on TSA plates before being transferred to TSA plates with or without antibiotics added at the indicated concentrations (created with BioRender.com). **(B)** Biofilms were homogenized after 24 and 48 hours of antibiotic exposure and plated to determine CFUs remaining versus immediately prior to antibiotic exposure. Data represent technical replicates of biological triplicates. The corresponding reduction in CFUs for a planktonic culture (mid-exponential phase) that was exposed to the same concentration of antibiotics for 24 hours is also superimposed. The difference in the reduction in CFUs between planktonic and biofilm cultures after 24 hours of antibiotic exposure is shown. 2-way ANOVA with Šídák multiple comparisons test; ****=$p$<0.0001.

**Figure S2**

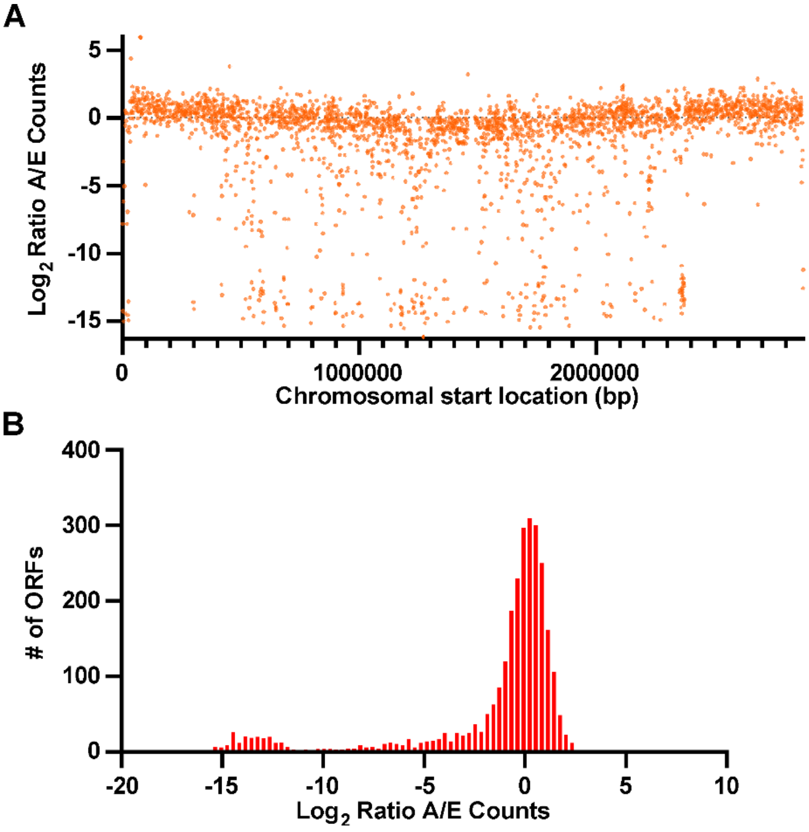


**Figure S2. Analysis of TnSeq library reveals high density library. (A)** $Log_2$ transformed ratios of the actual number of sequencing reads per gene/expected number of sequencing reads (A/E) is shown for each annotated open reading frame across the *S. aureus* USA300 FPR_3757 genome (A/E ratio). (**B**) Histogram showing the distribution of the $Log_2$ transformed A/E counts for all 2807 open reading frames annotated in the *S. aureus* USA300 FPR_3757 genome.

**Figure S3**

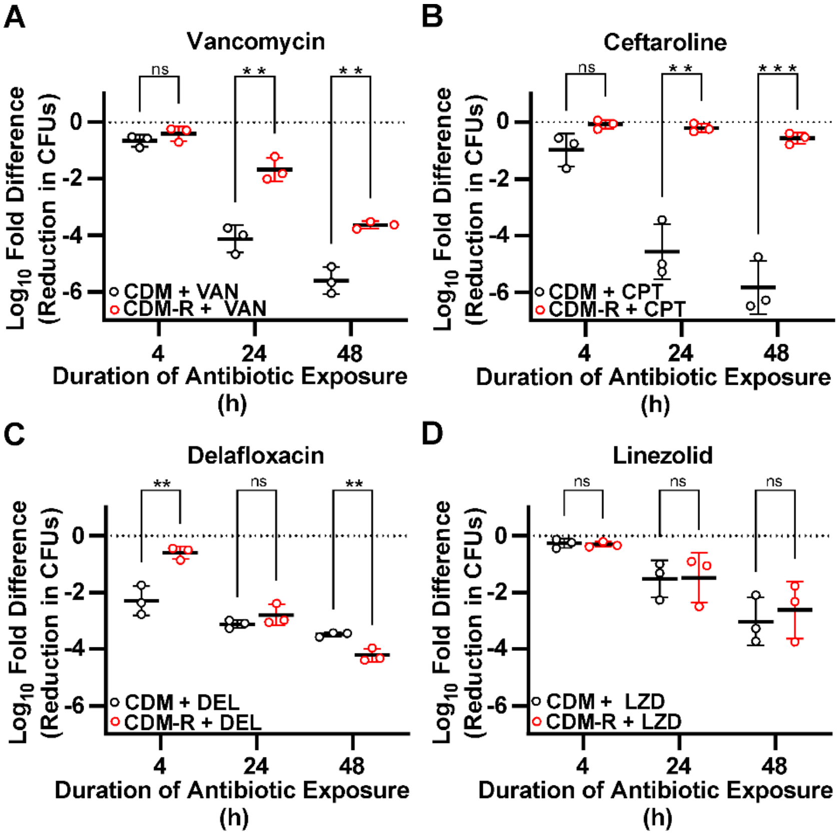


**Figure S3. Induction of antibiotic tolerance is more variable during planktonic growth.** Overnight cultures of JE2 were diluted 1:100 in CDM broth and grown to mid-log phase before being washed, split into fresh CDM or CDM-R broth with or without the addition of the indicated antibiotics, and grown for an additional 48 hours under planktonic growth conditions. Data represent technical replicates of biological triplicates. Student T-test; *=$p$<0.05, **=$p$<0.005, ***=$p$<0.0005, ****=$p$<0.0001, ns=not significant, VAN=vancomycin (400 µg/ml), CPT=ceftaroline (20 µg/ml), DEL=delafloxacin (9 µg/ml), LZD=linezolid (20 µg/ml). Data represent technical replicates of biological triplicates.

**Figure S4**

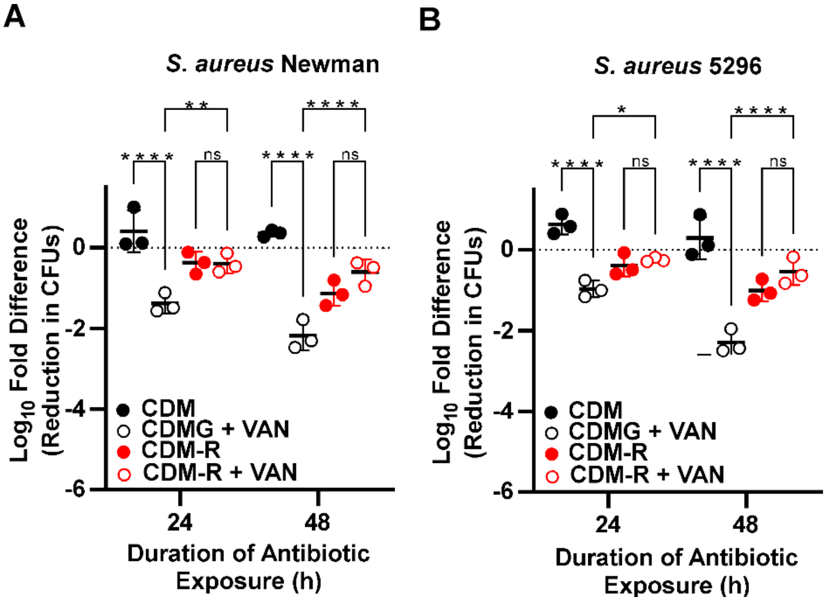


**Figure S4. Arginine deprivation induces tolerance in multiple strains of S. aureus.** The MSSA strain Newman (**A**) and the MRSA clinical isolate 5296 (**B**) were grown for 48 hours on polycarbonate filters on CDM agar plates before being homogenized and transferred to liquid CDM or CDM-R media with or without vancomycin added at a concentration of 400 µg/ml. Data represent technical replicates of biological triplicates. 2-way ANOVA with Tukey multiple comparisons test; *=$p$<0.05, **=$p$<0.005, ***=$p$<0.0005, ****=$p$<0.0001, ns=not significant, VAN=vancomycin.

**Figure S5**

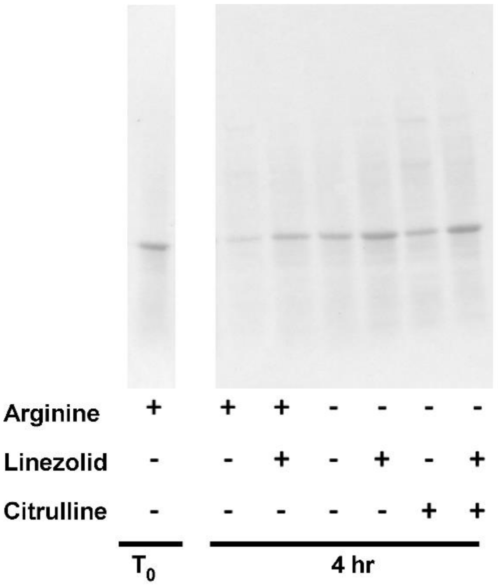


**Figure S5. Ponceau S Stain of Total Protein.** Total protein was isolated from homogenized biofilm cultures at $T_0$ and 4 hrs after incubation with the methionine analog, L-HPG. L-HPG incorporated into nascent protein was labelled with biotin via click chemistry and the resulting protein samples were separated via SDS-PAGE gel. Protein was transferred to a nitrocellulose membrane and prior to western blotting total protein was visualized using a Ponceau S stain. A representative Ponceau S stained membrane is shown here.

**Figure S6**

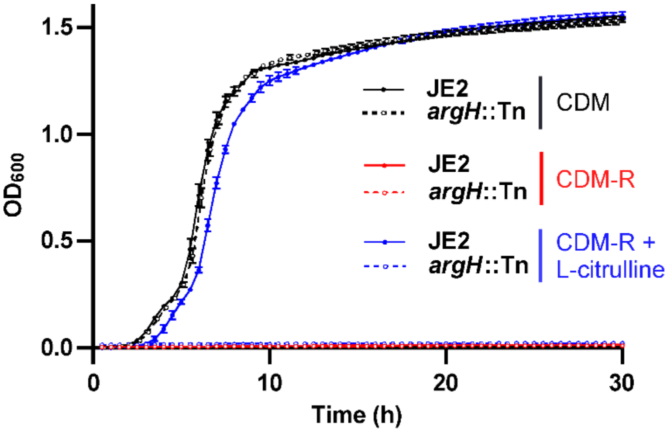


**Figure S6. ArgH is required to utilize citrulline for growth in the absence of arginine.** Growth of JE2 is inhibited in CDM in the absence of arginine, but this auxotrophy can be chemically complemented with the addition of citrulline. An *argH*::Tn mutant, however, is unable to use citrulline to complement an arginine auxotrophy.

**Figure S7**

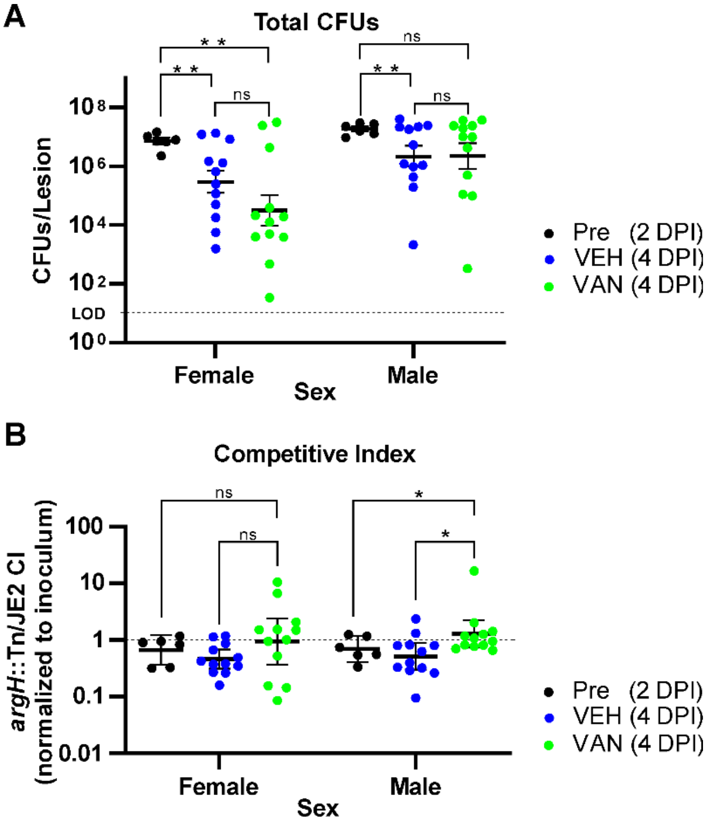


**Figure S7. Differences in the response to vancomycin treatment in a murine SSTI model by sex.** Both male and female C57BL/6 mice were infected with a mixture of JE2 and *argH*::Tn using a superficial skin infection model of a SSTI. Infected female mice exhibited significant decreases in CFUs between the 2 and 4 DPI, regardless of treatment group, while males only showed a significant decrease in CFUs between 2 and 4 DPI for the vehicle control group (**A**). Changes in the competitive index (CI) were similar between the two sexes (**B**) with both males and females showing a numerical increase in the CI to favor the mutant during vancomycin treatment, and this difference was significant for males. Data are combined from two independent experiments with a total of 6 mice per sex at 2 DPI and 12 mice per sex for each of the 4 DPI treatment groups. (**A**) Multiple T-test with Holm- Šídák multiple comparisons test, (**B**) Mann-Whitney test; *=p<0.05, **=p<0.005, ns=not significant, VEH= no antibiotic vehicle control, VAN=vancomycin, DPI=days post infection.

**Table S1.** ***S. aureus*** **proteins identified as having a significant (adj. p value ≤0.05, z-score of $log_2$ fold change ≥1 or ≤-1) difference in abundance in the presence of one or more antibiotics during biofilm growth.**

| Proteins with significant increase in abundance in the presence of antibiotics during biofilm growth | | | | | | |
|---|---|---|---|---|---|---|
| | | | Log2 Fold Change (vs untreated) | | | |
| Gene_ID | Gene Name | Gene Product | Vancomycin | Ceftaroline | Delafloxacin | Linezolid |
| SAUSA300_RS00330 | *arcA.2* | arginine deiminase | 1.25 | | | |
| SAUSA300_RS01260 | *glcC* | PTS glucose transporter subunit IIB | | | | 1.17 |
| SAUSA300_RS01920 | - | mechanosensitive ion channel protein | 1.31 | 1.88 | | |
| SAUSA300_RS01930 | *ychF* | GTP-binding protein | | 1.35 | | |
| SAUSA300_RS02105 | - | NAD(P)-dependent oxidoreductase | | | | 3.10 |
| SAUSA300_RS02565 | *prs* | ribose-phosphate pyrophosphokinase | | 1.00 | | |
| SAUSA300_RS02875 | - | UDP-glucose 4-epimerase | | 1.22 | | |
| SAUSA300_RS02960 | *hxlA* | 3-hexulose-6-phosphate synthase | | 1.17 | | |
| SAUSA300_RS03420 | *dhaM* | PTS-dependent dihydroxyacetone kinase phosphotransferase subunit | | 2.27 | | |
| SAUSA300_RS03450 | - | N-acetyltransferase | 1.31 | | | |
| SAUSA300_RS03715 | *saeQ* | hypothetical protein | | | | 1.00 |
| SAUSA300_RS03845 | *nrdE* | ribonucleotide-diphosphate reductase subunit alpha | | | | 1.49 |
| SAUSA300_RS03850 | *nrdF* | ribonucleotide-diphosphate reductase subunit beta | | | | 1.32 |
| SAUSA300_RS04000 | *uvrB* | excinuclease ABC subunit B | | | | 1.44 |
| SAUSA300_RS04005 | *uvrA* | excinuclease ABC subunit A | | | | 1.99 |
| SAUSA300_RS04380 | - | phage capsid protein | | | | 2.65 |
| SAUSA300_RS04560 | *ndh2* | NADH dehydrogenase | 1.05 | | | |
| SAUSA300_RS04680 | *spsA* | inactive signal peptidase IA | | 2.05 | | |
| SAUSA300_RS04695 | *addA* | ATP-dependent helicase/nuclease subunit A | | 1.14 | | |
| SAUSA300_RS04710 | *cdr* | CoA-disulfide reductase | | 2.09 | | |
| SAUSA300_RS04730 | *clpB* | chaperone protein | | 1.66 | | |
| SAUSA300_RS04905 | *fabI* | enoyl-ACP reductase | | 1.05 | | |
| SAUSA300_RS04995 | *lplA1* | lipoate--protein ligase A | | 1.14 | | |
| SAUSA300_RS05155 | *fmtA* | teichoic acid D-Ala esterase | | 1.51 | | |
| SAUSA300_RS05195 | *purE* | 5-(carboxyamino)imidazole ribonucleotide mutase | | 3.64 | | |
| SAUSA300_RS05305 | *cydA* | cytochrome ubiquinol oxidase subunit I | | 1.34 | | |
| SAUSA300_RS05385 | *potB* | spermidine/putrescine ABC transporter permease | | | | 2.36 |

| | | | | | | |
|---|---|---|---|---|---|---|
| SAUSA300_RS05830 | *pbp1* | penicillin-binding protein | | 1.69 | | |
| SAUSA300_RS05875 | - | cell division protein | | | 1.28 | |
| SAUSA300_RS06235 | *pyrH* | UMP kinase | | 1.00 | | 1.47 |
| SAUSA300_RS06245 | *uppS* | isoprenyl transferase | | 1.90 | | |
| SAUSA300_RS06260 | *proS* | proline--tRNA ligase | | 1.54 | | |
| SAUSA300_RS06370 | *recA* | DNA recombination/repair protein | | | | 3.26 |
| SAUSA300_RS06695 | *guaC* | guanosine monophosphate reductase | | 1.14 | | |
| SAUSA300_RS06745 | *sbcD* | exonuclease sbcCD subunit D | | | | 2.48 |
| SAUSA300_RS06945 | *phoU* | phosphate transport system regulatory protein | | | | 7.47 |
| SAUSA300_RS06965 | *pstS* | phosphate-binding protein | 2.14 | | | |
| SAUSA300_RS07165 | *crr* | glucose-specific phosphotransferase enzyme IIA component | 1.43 | 1.71 | | |
| SAUSA300_RS07170 | *msrB* | peptide-methionine (R)-S-oxide reductase | | 1.62 | | |
| SAUSA300_RS07175 | *msrA1* | peptide-methionine (S)-S-oxide reductase | | 1.35 | | |
| SAUSA300_RS07315 | *pbp2* | penicillin-binding protein 2 | | 1.63 | | |
| SAUSA300_RS08000 | *bfmBAB* | alpha-ketoacid dehydrogenase subunit beta | | 1.19 | | |
| SAUSA300_RS08175 | *gcvT* | aminomethyltransferase | | 1.07 | | |
| SAUSA300_RS08750 | - | DUF4930 domain-containing protein | 3.30 | 3.71 | | 1.69 |
| SAUSA300_RS09130 | *ptaA* | PTS glucose transporter subunit IIBC | | | 1.29 | |
| SAUSA300_RS09140 | *htrA1* | serine protease | 1.45 | 1.66 | | |
| SAUSA300_RS09800 | *prsA* | foldase | 1.54 | | | |
| SAUSA300_RS10135 | *sgtB* | monofunctional glycosyltransferase | 2.49 | 2.88 | | |
| SAUSA300_RS10185 | *vraR* | DNA-binding response regulator | | 2.56 | | |
| SAUSA300_RS10190 | *vraS* | two-component sensor histidine kinase | 2.07 | 2.67 | | |
| SAUSA300_RS10195 | *vraT* | transporter | | 2.00 | | |
| SAUSA300_RS10245 | *murT* | UDP-N-acetylmuramate--alanine ligase | | 1.50 | | 1.08 |
| SAUSA300_RS10540 | *sak* | staphylokinase | | | | 5.67 |
| SAUSA300_RS10570 | - | Phi77 ORF044-like protein | | | | 2.68 |
| SAUSA300_RS10590 | - | Immunodominant staphylococcal antigen A | | | | 2.50 |
| SAUSA300_RS10600 | - | Conserved hypothetical phage protein | | | | 5.31 |
| SAUSA300_RS10605 | - | Phi77 ORF020-like protein, phage major tail protein | | 1.57 | | 6.62 |
| SAUSA300_RS10630 | - | Phi77 ORF045-like protein | | | | 3.37 |
| SAUSA300_RS10635 | - | Phi77 ORF006-like protein, putative capsid protein | | | | 8.22 |
| SAUSA300_RS10645 | - | Phage portal protein | | | | 3.16 |
| SAUSA300_RS10680 | - | Phi77 ORF031-like protein | | | | 3.62 |
| SAUSA300_RS10700 | - | Conserved hypothetical phage protein | | | | 6.07 |
| SAUSA300_RS10750 | *recT* | Putative phage-related DNA recombination protein | | | | 8.03 |

| | | | | | | |
|---|---|---|---|---|---|---|
| SAUSA300_RS10775 | - | Conserved hypothetical phage protein | | | | 5.67 |
| SAUSA300_RS10785 | - | Phi77 ORF014-like protein, phage anti-repressor protein | | | | 4.57 |
| SAUSA300_RS11220 | *ddl* | D-alanine--D-alanine ligase | | 1.24 | | |
| SAUSA300_RS11430 | - | aldehyde dehydrogenase family protein | | | 1.61 | |
| SAUSA300_RS11440 | *murA2* | UDP-N-acetylglucosamine 1-carboxyvinyltransferase | | 1.43 | | |
| SAUSA300_RS11720 | *sdrM* | MFS transporter | | | | 1.37 |
| SAUSA300_RS11835 | - | alpha/beta hydrolase | | 1.33 | | |
| SAUSA300_RS12480 | - | transcriptional regulator | 1.64 | 2.32 | | |
| SAUSA300_RS12720 | *tcaA* | zinc ribbon domain-containing protein | 1.36 | 1.69 | | |
| SAUSA300_RS13070 | *hlgA* | gamma-hemolysin component A | | 2.57 | | |
| SAUSA300_RS13155 | - | membrane protein | 3.68 | 3.98 | | |
| SAUSA300_RS13210 | - | NAD(P)-dependent oxidoreductase | 1.12 | | | |
| SAUSA300_RS13270 | *pnbA* | carboxylesterase/lipase family protein | | 1.15 | | |
| SAUSA300_RS13560 | - | MerR family transcriptional regulator | | 2.77 | | |
| SAUSA300_RS13565 | *relP* | GTP pyrophosphokinase | 2.69 | | | |
| SAUSA300_RS14290 | *arcB.1* | ornithine carbamoyltransferase | | | | 1.17 |
| SAUSA300_RS14295 | *arcA.1* | arginine deiminase | | | | 1.35 |
| SAUSA300_RS14450 | *icaR* | biofilm operon icaADBC HTH-type negative transcriptional regulator | | 3.27 | | 3.41 |
| SAUSA300_RS14475 | *gehA* | Lipase 1 | 1.22 | | | |
| SAUSA300_RS14565 | *drp35* | lactonase | 3.05 | 3.83 | | |
| SAUSA300_RS14615 | - | arylamine N-acetyltransferase | | 1.37 | | |

| Proteins with significant decrease in abundance in the presence of antibiotics during biofilm growth | | | | | | |
|---|---|---|---|---|---|---|
| | | | Log2 Fold Change (vs untreated) | | | |
| Gene_ID | Gene Name | Gene Product | Vancomycin | Ceftaroline | Delafloxacin | Linezolid |
| SAUSA300_RS00110 | *walK* | cell wall metabolism sensor histidine kinase | -2.51 | | | |
| SAUSA300_RS00590 | *sarS* | transcriptional regulator | | -1.21 | | |
| SAUSA300_RS00735 | *deoC1* | 2-deoxyribose-5-phosphate aldolase | -2.60 | | | |
| SAUSA300_RS01490 | *esxA* | Type VII secretion system extracellular protein A | | -1.93 | | -2.58 |
| SAUSA300_RS01985 | - | transcriptional regulator | | -1.34 | | -1.94 |
| SAUSA300_RS02905 | - | HAD family phosphatase | -1.03 | | | |
| SAUSA300_RS02920 | *sdrD* | serine-aspartate repeat-containing protein D | | -1.05 | | |

| | | | | | | |
|---|---|---|---|---|---|---|
| SAUSA300_RS02970 | - | HAD family hydrolase | | -1.03 | | |
| SAUSA300_RS03765 | - | allophanate hydrolase | | | | -1.65 |
| SAUSA300_RS04335 | - | transcriptional regulator | | -1.14 | | |
| SAUSA300_RS04475 | *lipA* | lipoyl synthase | | -1.31 | | |
| SAUSA300_RS04535 | *nfu* | NifU family protein | | -1.87 | | -2.09 |
| SAUSA300_RS04655 | *glpQ* | glycerophosphodiester phosphodiesterase | | -1.33 | | |
| SAUSA300_RS04660 | *argH* | argininosuccinate lyase | -2.29 | -2.89 | | |
| SAUSA300_RS04665 | *argG* | argininosuccinate synthase | -2.23 | -2.78 | | |
| SAUSA300_RS04805 | *opp-3A* | peptide ABC transporter substrate-binding protein | | -1.37 | | |
| SAUSA300_RS04840 | *spxA* | regulatory protein | | | | -1.50 |
| SAUSA300_RS05000 | - | DUF2187 domain-containing protein | -2.61 | | | |
| SAUSA300_RS05040 | - | hypothetical protein | | -1.42 | | |
| SAUSA300_RS05135 | *atl* | bifunctional autolysin | -1.10 | | | |
| SAUSA300_RS05540 | *isdA* | iron-regulated surface determinant protein A | | -1.14 | | -1.64 |
| SAUSA300_RS05760 | *arcC.3* | carbamate kinase | -2.04 | | | |
| SAUSA300_RS06340 | - | peptidase M16 | | -1.69 | | |
| SAUSA300_RS06380 | - | hypothetical protein | | -1.17 | | |
| SAUSA300_RS06590 | - | hypothetical protein | | | | -1.04 |
| SAUSA300_RS06645 | *hom* | homoserine dehydrogenase | -1.05 | | | |
| SAUSA300_RS06650 | *thrC* | threonine synthase | | | | -1.57 |
| SAUSA300_RS06680 | *katA* | catalase | | | | -1.92 |
| SAUSA300_RS06685 | *rpmG2* | 50S ribosomal protein L33 | -7.38 | | | |
| SAUSA300_RS06995 | *asd* | aspartate-semialdehyde dehydrogenase | | -1.46 | | -1.74 |
| SAUSA300_RS07170 | *msrB* | peptide-methionine (R)-S-oxide reductase | | | | -1.14 |
| SAUSA300_RS07195 | - | hypothetical protein | -1.06 | | | |
| SAUSA300_RS07320 | - | hypothetical protein | -2.22 | | | |
| SAUSA300_RS07825 | - | transcriptional regulator | | -1.26 | | |
| SAUSA300_RS08310 | - | phosphoenolpyruvate synthetase regulatory protein | -1.10 | | | |
| SAUSA300_RS08375 | *mtaB* | tRNA (N(6)-L-threonylcarbamoyladenosine(37)-C(2))- methylthiotransferase | -2.22 | -1.90 | -1.08 | |
| SAUSA300_RS08530 | - | acetyl-CoA carboxylase biotin carboxyl carrier protein subunit | | -1.12 | | -2.29 |
| SAUSA300_RS08620 | *csbD* | CsbD family protein | -4.57 | -1.41 | | |
| SAUSA300_RS08865 | *rplT* | 50S ribosomal protein L20 | | -1.01 | | |
| SAUSA300_RS08870 | *rpmI* | 50S ribosomal protein L35 | -6.25 | | | |
| SAUSA300_RS09465 | *metK* | S-adenosylmethionine synthase | -1.67 | | | |
| SAUSA300_RS09805 | *cbf1* | 3'-5' exoribonuclease | | | | -1.31 |
| SAUSA300_RS09825 | - | hypothetical protein | | | -1.08 | |

| | | | | | | |
|---|---|---|---|---|---|---|
| SAUSA300_RS09835 | - | transcriptional regulator | | -1.22 | | |
| SAUSA300_RS09840 | *airR* | DNA-binding response regulator | | | | -2.91 |
| SAUSA300_RS09890 | *glnQ* | amino acid ABC transporter ATP-binding protein | -1.53 | -3.33 | | |
| SAUSA300_RS10060 | *perR* | transcriptional repressor | | -1.80 | | |
| SAUSA300_RS10175 | - | hypothetical protein | -1.41 | -2.11 | | |
| SAUSA300_RS11270 | *thiE* | thiamine phosphate synthase | -1.86 | | | |
| SAUSA300_RS11705 | *salA* | chromosome partitioning protein | -1.98 | | | |
| SAUSA300_RS12060 | *rpmD* | 50S ribosomal protein L30 | | -1.05 | | |
| SAUSA300_RS12270 | *moaD* | molybdopterin synthase sulfur carrier subunit | | -2.79 | | -1.79 |
| SAUSA300_RS12290 | *moaC* | cyclic pyranopterin monophosphate synthase | | -1.10 | | |
| SAUSA300_RS12390 | *sarR* | transcriptional regulator | | | | -1.32 |
| SAUSA300_RS13570 | - | hypothetical protein | | -2.89 | | -2.50 |
| SAUSA300_RS13860 | *copZ* | copper chaperone | | -1.80 | | |
| SAUSA300_RS13915 | *isaA* | transglycosylase IsaA | | -1.50 | | |
| SAUSA300_RS14025 | - | hypothetical protein | | -1.52 | | |
| SAUSA300_RS14280 | *arc.1* | carbamate kinase | -1.27 | | | |

**Table S2.** ***S. aureus*** **genes identified via TnSeq as having a significant (adj. p value ≤0.05, z-score of $log_2$ fold change ≥1 or ≤-1) effect on fitness in the presence of one or more antibiotics during biofilm growth.**

| Genes with significant decrease in fitness in the presence of antibiotics due to transposon insertions | | | | | | |
|---|---|---|---|---|---|---|
| | | | Log2 Fold Change (vs untreated) | | | |
| Gene_ID | Gene Name | Gene Product | Vancomycin | Ceftaroline | Delafloxacin | Linezolid |
| SAUSA300_RS00260 | - | hypothetical protein | -1.55 | | | |
| SAUSA300_RS00715 | - | GntR family transcriptional regulator | | | | -1.65 |
| SAUSA300_RS01375 | *bglR* | GntR family transcriptional regulator | -1.17 | | | |
| SAUSA300_RS01560 | - | hypothetical protein | -2.39 | | | |
| SAUSA300_RS02535 | *ipk* | 4-diphosphocytidyl-2C-methyl-D-erythritol kinase | -1.03 | | | -1.29 |
| SAUSA300_RS02540 | *purR* | pur operon repressor | -2.38 | -1.35 | -1.15 | -2.03 |
| SAUSA300_RS02545 | *yabJ* | RidA family protein | -5.38 | | | |
| SAUSA300_RS02550 | *spoVG* | stage V sporulation protein G | -2.58 | | | |
| SAUSA300_RS02950 | - | hypothetical protein | -2.71 | | | |
| SAUSA300_RS03085 | - | transcriptional regulator | -1.44 | | | |
| SAUSA300_RS03300 | *mnhF2* | cation:proton antiporter | -2.88 | | | |
| SAUSA300_RS03370 | *pbp4* | D-alanyl-D-alanine carboxypeptidase | -1.59 | | | |
| SAUSA300_RS03455 | *graX* | NAD(P)-dependent oxidoreductase | -4.31 | | | |
| SAUSA300_RS03460 | *graR* | DNA-binding response regulator | -3.16 | | | |
| SAUSA300_RS03465 | *graS* | sensor histidine kinase | -4.26 | | | |
| SAUSA300_RS03470 | *vraF* | ABC transporter ATP-binding protein | -3.82 | -1.05 | | |
| SAUSA300_RS03475 | *vraG* | ABC transporter permease | -3.71 | -1.07 | | |
| SAUSA300_RS03605 | *mgrA* | MarR family transcriptional regulator | | -1.12 | | |
| SAUSA300_RS03790 | *recQ1* | DNA helicase | -1.18 | | | |
| SAUSA300_RS03820 | - | hypothetical protein | -2.19 | | | |
| SAUSA300_RS03930 | *gdpS* | GGDEF domain-containing protein | -3.07 | -2.00 | | |
| SAUSA300_RS04355 | - | hypothetical protein | -1.54 | | | |
| SAUSA300_RS04570 | - | sodium:proton antiporter | -1.60 | | | |
| SAUSA300_RS04930 | *ltaA* | MFS transporter | | | | -1.95 |
| SAUSA300_RS05025 | - | hypothetical protein | -1.59 | | | |
| SAUSA300_RS05085 | *menD* | 2-succinyl-5-enolpyruvyl-6-hydroxy-3- cyclohexene-1-carboxylate synthase | -1.90 | | | |
| SAUSA300_RS05155 | *fmtA* | teichoic acid D-Ala esterase | -1.40 | | | |
| SAUSA300_RS05255 | *thiW* | ABC transporter ATP-binding protein | -1.29 | | | |

| | | | | | | |
|---|---|---|---|---|---|---|
| SAUSA300_RS05680 | *flr* | FPRL1 inhibitory protein | -2.77 | | | |
| SAUSA300_RS06055 | *fakA* | fatty acid kinase catalytic subunit | | | | -1.07 |
| SAUSA300_RS06185 | *topA* | DNA topoisomerase I | -2.32 | | | |
| SAUSA300_RS06200 | *hslV* | HslU--HslV peptidase proteolytic subunit | | | -1.76 | -2.09 |
| SAUSA300_RS06205 | *hslU* | HslU--HslV peptidase ATPase subunit | | | -1.13 | -1.09 |
| SAUSA300_RS06350 | - | hypothetical protein | -1.45 | | | |
| SAUSA300_RS06525 | - | hypothetical protein | -2.28 | | | |
| SAUSA300_RS06820 | *mprF* | phosphatidylglycerol lysyltransferase | -3.50 | | | |
| SAUSA300_RS07110 | *arlS* | two-component sensor histidine kinase | -1.65 | | | |
| SAUSA300_RS07115 | *arlR* | DNA-binding response regulator | -2.72 | | | |
| SAUSA300_RS07195 | - | BrxA/BrxB family bacilliredoxin | | | -1.62 | -1.69 |
| SAUSA300_RS07300 | - | YppE family protein | -2.18 | | | |
| SAUSA300_RS07400 | - | hypothetical protein | -2.19 | | | |
| SAUSA300_RS07765 | - | DNA polymerase | -1.15 | | | |
| SAUSA300_RS07850 | - | hypothetical protein | -2.14 | | | -1.30 |
| SAUSA300_RS08375 | *mtaB* | tRNA (N(6)-L-threonylcarbamoyladenosine(37)-C(2))-methylthiotransferase | -1.69 | | | |
| SAUSA300_RS09085 | - | GAF domain-containing protein | -2.81 | | | |
| SAUSA300_RS09365 | *ribB* | riboflavin synthase | -1.41 | | | |
| SAUSA300_RS09830 | - | DUF445 domain-containing protein | -1.19 | | | |
| SAUSA300_RS09850 | - | RluA family pseudouridine synthase | -1.85 | | | |
| SAUSA300_RS10435 | - | DUF4097 family beta strand repeat-containing protein | -1.07 | | | |
| SAUSA300_RS10440 | - | DUF1700 domain-containing protein | -1.20 | | | |
| SAUSA300_RS10445 | - | membrane protein | -1.63 | | | |
| SAUSA300_RS10635 | - | phage major capsid protein | -1.40 | | | |
| SAUSA300_RS10700 | - | hypothetical protein | -3.37 | | | |
| SAUSA300_RS10705 | - | DUF1024 family protein | -6.93 | | | |
| SAUSA300_RS10935 | *agrB* | accessory gene regulator | -2.24 | | | -1.07 |
| SAUSA300_RS10940 | *agrD* | cyclic lactone autoinducer peptide | -1.67 | | | -1.03 |
| SAUSA300_RS10945 | *agrC* | ATP-binding protein | -1.89 | | | -1.09 |
| SAUSA300_RS10950 | *agrA* | DNA-binding response regulator | -2.22 | | | -1.15 |
| SAUSA300_RS10965 | *scrB* | sucrose-6-phosphate hydrolase | -1.38 | | | -1.39 |
| SAUSA300_RS10970 | *scrR* | LacI family transcriptional regulator | -1.25 | | | |
| SAUSA300_RS11055 | *leuA* | 2-isopropylmalate synthase | -1.02 | | | |
| SAUSA300_RS11120 | *sigB* | RNA polymerase sigma factor | -3.35 | | | |
| SAUSA300_RS11125 | *rsbW* | anti-sigma B factor | -2.85 | | | |
| SAUSA300_RS11130 | *rsbV* | anti-sigma B factor antagonist | -3.42 | | | |
| SAUSA300_RS11135 | *rsbU* | serine phosphatase | -3.61 | | | -1.05 |
| SAUSA300_RS11145 | *mazF* | type II toxin-antitoxin system PemK/MazF family toxin | -2.26 | | | |

| | | | | | | |
|---|---|---|---|---|---|---|
| SAUSA300_RS11150 | - | type II toxin-antitoxin system antitoxin | -3.43 | | | |
| SAUSA300_RS11320 | - | membrane protein | -1.36 | | | |
| SAUSA300_RS11380 | *upp* | uracil phosphoribosyltransferase | -1.62 | -1.98 | -2.28 | -2.29 |
| SAUSA300_RS11385 | *glyA* | serine hydroxymethyltransferase | -1.52 | -1.18 | -1.57 | -1.76 |
| SAUSA300_RS11390 | - | TIGR01440 family protein | -1.60 | -1.35 | -1.52 | -1.29 |
| SAUSA300_RS11635 | *ybbR* | hypothetical protein | | -1.76 | -2.02 | -2.14 |
| SAUSA300_RS11725 | - | hemolysin III | | -1.49 | | |
| SAUSA300_RS11730 | - | uridylyltransferase | | -1.25 | | |
| SAUSA300_RS11935 | *budA1* | alpha-acetolactate decarboxylase | -1.24 | | | |
| SAUSA300_RS13300 | - | prevent-host-death protein | -1.58 | | | |
| SAUSA300_RS13325 | *cntE* | MFS transporter | | -1.26 | -1.04 | -1.98 |
| SAUSA300_RS14085 | *bcaP* | amino acid permease | -1.05 | | | |
| SAUSA300_RS14115 | - | AMP-binding protein | -1.59 | -1.75 | | -1.30 |
| SAUSA300_RS14125 | - | sterile alpha motif-like domain-containing protein | -1.93 | | | |
| SAUSA300_RS15875 | - | hypothetical protein | -2.26 | | | |
| SAUSA300_RS15890 | - | hypothetical protein | -2.23 | | | |

| Genes with significant increase in fitness in the presence of antibiotics due to transposon insertions | | | | | | |
|---|---|---|---|---|---|---|
| | | | Log2 Fold Change (vs untreated) | | | |
| Gene_ID | Gene Name | Gene Product | Vancomycin | Ceftaroline | Delafloxacin | Linezolid |
| SAUSA300_RS00090 | *purA* | adenylosuccinate synthetase | | 1.19 | | 1.51 |
| SAUSA300_RS00115 | *walH* | two-component system activity regulator | | 1.63 | 2.04 | 2.04 |
| SAUSA300_RS00120 | *walI* | two-component system regulatory protein | | 1.62 | 1.89 | 1.86 |
| SAUSA300_RS01015 | *murQ* | N-acetylmuramic acid 6-phosphate etherase | | 1.21 | 1.51 | |
| SAUSA300_RS01930 | *ychF* | GTP-binding protein | | | 1.28 | 1.39 |
| SAUSA300_RS02585 | *mfd* | transcription-repair coupling factor | | 1.69 | 1.46 | 1.12 |
| SAUSA300_RS02635 | *cysK* | cysteine synthase | | | | 1.08 |
| SAUSA300_RS03175 | - | N-acetyltransferase | | 1.30 | 1.08 | 1.25 |
| SAUSA300_RS03240 | - | hypothetical protein | | 1.32 | 1.63 | 1.37 |
| SAUSA300_RS03315 | *mntC* | metal ABC transporter substrate-binding protein | | 1.66 | 1.73 | 1.67 |
| SAUSA300_RS03320 | *mntB* | metal ABC transporter permease | | 1.97 | 1.75 | 1.76 |
| SAUSA300_RS03325 | *mntA* | phosphonate ABC transporter ATP-binding protein | | 1.40 | 1.24 | |
| SAUSA300_RS03525 | - | hypothetical protein | | 3.48 | 3.25 | 3.40 |
| SAUSA300_RS03530 | *ccpE* | LysR family transcriptional regulator | | 1.91 | | 1.99 |

| | | | | | | |
|---|---|---|---|---|---|---|
| SAUSA300_RS04270 | *gcvH* | glycine cleavage system protein H | | | 4.08 | |
| SAUSA300_RS04465 | - | anion permease | | | | 1.42 |
| SAUSA300_RS04560 | *ndh2* | NADH dehydrogenase | | 3.31 | 3.03 | 3.28 |
| SAUSA300_RS04660 | *argH* | argininosuccinate lyase | | 1.45 | 1.11 | 1.41 |
| SAUSA300_RS04795 | *opp-3D* | ABC transporter ATP-binding protein | | 1.94 | 2.05 | 1.75 |
| SAUSA300_RS05170 | *qoxB* | cytochrome ubiquinol oxidase subunit I | | 1.95 | | |
| SAUSA300_RS05200 | *purK* | 5-(carboxyamino)imidazole ribonucleotide synthase | | 1.15 | 1.22 | 1.56 |
| SAUSA300_RS05230 | *purM* | phosphoribosylformylglycinamidine cyclo-ligase | | | | 1.58 |
| SAUSA300_RS05240 | *purH* | bifunctional purine biosynthesis protein | | 1.16 | | 1.20 |
| SAUSA300_RS05245 | *purD* | phosphoribosylamine--glycine ligase | | 1.54 | 1.62 | 1.71 |
| SAUSA300_RS05460 | *ctaA* | heme A synthase | | 2.08 | | 2.61 |
| SAUSA300_RS05600 | *zapA* | cell division protein | | | 1.05 | |
| SAUSA300_RS05635 | *sdhC* | succinate dehydrogenase cytochrome B558 | | | 2.34 | 2.22 |
| SAUSA300_RS05640 | *sdhA* | succinate dehydrogenase flavoprotein subunit | | 1.27 | 1.35 | 1.78 |
| SAUSA300_RS05895 | - | glyoxalase | | | | 1.08 |
| SAUSA300_RS05925 | *pyrB* | aspartate carbamoyltransferase | | | 1.55 | |
| SAUSA300_RS05930 | *pyrC* | dihydroorotase | | 1.73 | 2.49 | 2.75 |
| SAUSA300_RS05935 | *carA* | carbamoyl-phosphate synthase small chain | | | | 1.79 |
| SAUSA300_RS05940 | *carB* | carbamoyl-phosphate synthase large chain | | 1.83 | 1.84 | 1.97 |
| SAUSA300_RS06445 | *glpD* | aerobic glycerol-3-phosphate dehydrogenase | | 1.63 | 1.61 | 1.84 |
| SAUSA300_RS06740 | - | membrane protein | | 2.07 | 2.14 | 1.66 |
| SAUSA300_RS06765 | *citB* | aconitate hydratase | | | 1.97 | |
| SAUSA300_RS07050 | *acyP* | acylphosphatase | | 1.43 | 1.39 | 1.48 |
| SAUSA300_RS07100 | *sucB* | dihydrolipoyllysine-residue succinyltransferase | | 2.26 | 2.51 | 2.42 |
| SAUSA300_RS07105 | *sucA* | 2-oxoglutarate dehydrogenase E1 component | | 1.55 | 1.45 | 1.91 |
| SAUSA300_RS07470 | *ansA* | L-asparaginase | | 1.10 | | |
| SAUSA300_RS08025 | *ispA* | geranyltranstransferase | | 2.76 | | |
| SAUSA300_RS08035 | *xseA* | exodeoxyribonuclease 7 large subunit | | 1.45 | | |
| SAUSA300_RS08460 | - | haloacid dehalogenase | | 1.68 | 1.95 | 1.63 |
| SAUSA300_RS08475 | *aroE* | shikimate dehydrogenase | | 3.13 | 2.89 | 2.68 |
| SAUSA300_RS08640 | - | hypothetical protein | 2.01 | | | |
| SAUSA300_RS08950 | *citC* | isocitrate dehydrogenase (NADP(+)) | | | 2.34 | |
| SAUSA300_RS09065 | *thiI* | tRNA 4-thiouridine(8) synthase | | | 1.27 | 1.58 |
| SAUSA300_RS09810 | - | DNA double-strand break repair Rad50 ATPase | | 1.13 | 1.52 | |
| SAUSA300_RS09895 | *glnP* | ABC transporter permease | 1.79 | | | |
| SAUSA300_RS10185 | *vraR* | DNA-binding response regulator | | | 3.65 | 3.22 |
| SAUSA300_RS10190 | *vraS* | two-component sensor histidine kinase | | | 3.45 | 4.11 |
| SAUSA300_RS10195 | *vraT* | cell wall-active antibiotics response protein | | | 1.86 | 1.99 |

| | | | | | | |
|---|---|---|---|---|---|---|
| SAUSA300_RS10335 | *purB* | adenylosuccinate lyase | | 1.60 | | 2.04 |
| SAUSA300_RS10450 | - | thioredoxin family protein | | | 1.19 | |
| SAUSA300_RS10465 | *pmtB* | ABC-2 transporter family protein | | 1.14 | | 1.27 |
| SAUSA300_RS10470 | *pmtA* | ABC transporter ATP-binding protein | | 1.74 | | |
| SAUSA300_RS10490 | *aspB* | aminotransferase class I/II-fold pyridoxal phosphate-dependent enzyme | | | | |
| SAUSA300_RS10990 | *rex* | transcriptional regulator | | 1.28 | | |
| SAUSA300_RS11210 | *cshA* | DEAD/DEAH box family ATP-dependent RNA helicase | | 1.76 | 2.11 | 1.97 |
| SAUSA300_RS11425 | *rho* | transcription termination factor Rho | | 1.49 | 1.39 | 1.77 |
| SAUSA300_RS11545 | *manA* | mannose-6-phosphate isomerase | 1.99 | | | |
| SAUSA300_RS11705 | *salA* | chromosome partitioning protein | | 1.62 | 1.45 | 2.17 |
| SAUSA300_RS11740 | - | hypothetical protein | | 2.10 | 1.74 | 2.24 |
| SAUSA300_RS11875 | *lacR* | DeoR/GlpR transcriptional regulator | 1.79 | 1.21 | | |
| SAUSA300_RS12165 | *pbuG* | NCS2 family permease | 1.86 | 1.16 | | |
| SAUSA300_RS12585 | *hutI* | imidazolonepropionase | | 1.12 | | 1.14 |
| SAUSA300_RS12610 | *lyrA* | lysostaphin resistance protein A | 1.72 | | | |
| SAUSA300_RS12630 | *galM* | galactose mutarotase | 1.12 | | | |
| SAUSA300_RS12770 | - | DUF3021 domain-containing protein | 1.48 | 1.41 | 1.05 | 1.40 |
| SAUSA300_RS12775 | *mqo* | malate:quinone oxidoreductase | | 1.32 | 2.03 | 1.85 |
| SAUSA300_RS12820 | - | DUF2871 domain-containing protein | | | | 1.01 |
| SAUSA300_RS12855 | *rsp* | transcriptional regulator | 1.46 | | | |
| SAUSA300_RS12865 | - | DUF4889 domain-containing protein | | | | 1.22 |
| SAUSA300_RS13840 | *rocA* | L-glutamate gamma-semialdehyde dehydrogenase | | | | 1.22 |
| SAUSA300_RS14135 | *betB* | betaine-aldehyde dehydrogenase | | | | 1.09 |
| SAUSA300_RS16025 | - | exotoxin | | | 1.10 | 1.00 |

**Table S3.** ***S. aureus*** **genes identified via TnSeq as having a significant (adj. p value ≤0.05, z-score of $\log_2$ fold change ≥1 or ≤-1) effect on fitness during biofilm growth.**

| Genes with significant decrease in fitness during biofilm growth due to transposon insertions | | | | |
|---|---|---|---|---|
| | | | $\text{Log}_2$ Fold Change (vs inoculum) | |
| Gene_ID | Gene Name | Gene Product | 48-hour Biofilm | 96-hour Biofilm |
| SAUSA300_RS00025 | *recF* | DNA replication and repair protein | -3.20 | -3.57 |
| SAUSA300_RS00090 | *purA* | adenylosuccinate synthetase | | -1.12 |
| SAUSA300_RS00115 | *walH* | hypothetical protein | -1.32 | -2.93 |
| SAUSA300_RS00120 | *walI* | hypothetical protein | | -2.00 |
| SAUSA300_RS00740 | *deoB* | phosphopentomutase | | -1.01 |
| SAUSA300_RS01015 | *murQ* | N-acetylmuramic acid 6-phosphate etherase | | -1.63 |
| SAUSA300_RS01930 | *ychF* | GTP-binding protein | -2.13 | -2.73 |
| SAUSA300_RS02045 | - | hypothetical protein | -1.22 | -1.64 |
| SAUSA300_RS02055 | - | hypothetical protein | | -1.13 |
| SAUSA300_RS02060 | *xpt* | xanthine phosphoribosyltransferase | | -1.47 |
| SAUSA300_RS02070 | *guaB* | IMP dehydrogenase | -2.18 | -3.02 |
| SAUSA300_RS02075 | *guaA* | GMP synthase (glutamine-hydrolyzing) | -3.28 | -2.15 |
| SAUSA300_RS02280 | *mpsA* | NADH dehydrogenase subunit 5 | -1.96 | -1.86 |
| SAUSA300_RS02315 | - | sodium-dependent transporter | -2.17 | -2.32 |
| SAUSA300_RS02400 | *treC* | glucohydrolase | | -1.65 |
| SAUSA300_RS02430 | *recR* | recombination protein | -3.00 | -2.87 |
| SAUSA300_RS02465 | *darA/pstA* | hypothetical protein | -4.29 | -4.26 |
| SAUSA300_RS02520 | *rnmV* | ribonuclease M5 | -1.60 | |
| SAUSA300_RS02525 | *ksgA* | ribosomal RNA small subunit methyltransferase A | -1.07 | -1.69 |
| SAUSA300_RS02585 | *mfd* | transcription-repair coupling factor | | -1.96 |
| SAUSA300_RS02610 | - | RNA-binding protein S1 | -3.21 | -3.18 |
| SAUSA300_RS02625 | *ftsH* | zinc metalloprotease | | -2.62 |
| SAUSA300_RS02635 | *cysK* | cysteine synthase | | -1.29 |
| SAUSA300_RS02700 | *pdxS* | pyridoxal 5'-phosphate synthase lyase subunit | -1.37 | -1.95 |
| SAUSA300_RS02730 | *clpC* | ATP-dependent Clp protease ATP-binding subunit | -1.21 | -1.17 |
| SAUSA300_RS02880 | *ilvE* | branched chain amino acid aminotransferase | -4.44 | -2.81 |
| SAUSA300_RS02900 | *tadA* | tRNA-specific adenosine deaminase | -4.38 | -4.19 |
| SAUSA300_RS02910 | *azo1* | FMN-dependent NADPH-azoreductase | | -1.18 |
| SAUSA300_RS03055 | *lipL* | biotin/lipoate A/B protein ligase family protein | -3.83 | -2.19 |
| SAUSA300_RS03240 | - | hypothetical protein | -1.01 | -2.30 |

| | | | | |
|---|---|---|---|---|
| SAUSA300_RS03250 | *sarA* | transcriptional regulator | -1.97 | -3.66 |
| SAUSA300_RS03315 | *mntC* | metal ABC transporter substrate-binding protein | -1.90 | -2.98 |
| SAUSA300_RS03320 | *mntB* | metal ABC transporter permease | -1.24 | -2.52 |
| SAUSA300_RS03325 | *mntA* | phosphonate ABC transporter ATP-binding protein | -1.18 | -1.92 |
| SAUSA300_RS03375 | *abcA* | ABC transporter ATP-binding protein | -1.21 | |
| SAUSA300_RS03480 | *pitR* | hypothetical protein | | -1.19 |
| SAUSA300_RS03525 | - | hypothetical protein | -1.14 | -4.03 |
| SAUSA300_RS03530 | *ccpE* | LysR family transcriptional regulator | | -1.75 |
| SAUSA300_RS03690 | *mpfA* | HlyC/CorC family transporter | | -1.29 |
| SAUSA300_RS03775 | *ltaS* | lipoteichoic acid synthase | | -1.50 |
| SAUSA300_RS03930 | *gdpS* | GGDEF domain-containing protein | -2.32 | |
| SAUSA300_RS03935 | *tagO* | undecaprenyl-phosphate alpha-N-acetylglucosaminyl 1-phosphate transferase | -1.16 | -1.87 |
| SAUSA300_RS04010 | *hprK* | HPr kinase/phosphorylase | -3.12 | -3.09 |
| SAUSA300_RS04015 | *lgt* | prolipoprotein diacylglyceryl transferase | | -1.39 |
| SAUSA300_RS04050 | *whiA* | sporulation regulator | | -2.75 |
| SAUSA300_RS04075 | *gapR* | transcriptional regulator | -2.02 | -1.99 |
| SAUSA300_RS04095 | *pgm* | 2,3-bisphosphoglycerate-independent phosphoglycerate mutase | -2.13 | -2.31 |
| SAUSA300_RS04110 | *secG* | protein-export membrane protein | | -2.38 |
| SAUSA300_RS04120 | *rnr* | ribonuclease R | -1.20 | -1.28 |
| SAUSA300_RS04225 | - | hypothetical protein | | -1.09 |
| SAUSA300_RS04250 | *aroD* | 3-dehydroquinase | | -1.82 |
| SAUSA300_RS04270 | *gcvH* | glycine cleavage system protein H | | -3.93 |
| SAUSA300_RS04335 | - | transcriptional regulator | -3.61 | -3.59 |
| SAUSA300_RS04465 | - | anion permease | | -1.31 |
| SAUSA300_RS04515 | *dltA* | D-alanine--poly(phosphoribitol) ligase | -1.38 | -1.35 |
| SAUSA300_RS04520 | *dltB* | D-alanyl-lipoteichoic acid biosynthesis protein | -4.86 | -4.82 |
| SAUSA300_RS04530 | *dltD* | D-alanyl-lipoteichoic acid biosynthesis protein | -1.59 | |
| SAUSA300_RS04535 | *nfu* | NifU family protein | -1.39 | -1.77 |
| SAUSA300_RS04550 | - | hypothetical protein | -2.16 | |
| SAUSA300_RS04560 | *ndh2* | NADH dehydrogenase | -1.23 | -3.52 |
| SAUSA300_RS04570 | - | sodium:proton antiporter | -1.14 | |
| SAUSA300_RS04645 | *gudB* | NAD-specific glutamate dehydrogenase | -1.94 | -1.72 |
| SAUSA300_RS04680 | *spsA* | inactive signal peptidase IA | | -1.68 |
| SAUSA300_RS04690 | *addB* | ATP-dependent helicase/deoxyribonuclease subunit B | -3.07 | -3.52 |
| SAUSA300_RS04695 | *addA* | ATP-dependent helicase/nuclease subunit A | -3.82 | -3.23 |
| SAUSA300_RS04795 | *opp-3D* | ABC transporter ATP-binding protein | | -1.53 |
| SAUSA300_RS04860 | *yjbH* | hypothetical protein | -3.05 | -4.39 |
| SAUSA300_RS04865 | *yjbI* | hypothetical protein | -1.80 | |

| | | | | |
|---|---|---|---|---|
| SAUSA300_RS04875 | - | hypothetical protein | -2.13 | -5.06 |
| SAUSA300_RS04880 | *relQ* | GTP pyrophosphokinase | -2.89 | -2.78 |
| SAUSA300_RS04930 | *ltaA* | MFS transporter | -3.23 | -2.25 |
| SAUSA300_RS04935 | *ugtP* | diacylglycerol beta-glucosyltransferase | -5.52 | -5.43 |
| SAUSA300_RS05095 | *menB* | 1,4-dihydroxy-2-naphthoyl-CoA synthase | -1.83 | -1.41 |
| SAUSA300_RS05160 | *qoxD* | quinol oxidase subunit 4 | | -3.20 |
| SAUSA300_RS05165 | *qoxC* | quinol oxidase subunit 3 | -1.69 | -4.43 |
| SAUSA300_RS05170 | *qoxB* | cytochrome ubiquinol oxidase subunit I | -3.24 | -3.07 |
| SAUSA300_RS05175 | *qoxA* | quinol oxidase subunit 2 | -2.79 | -2.90 |
| SAUSA300_RS05195 | *purE* | 5-(carboxyamino)imidazole ribonucleotide mutase | | -2.20 |
| SAUSA300_RS05200 | *purK* | 5-(carboxyamino)imidazole ribonucleotide synthase | | -1.41 |
| SAUSA300_RS05205 | *purC* | phosphoribosylaminoimidazolesuccinocarboxamide synthase | | -1.21 |
| SAUSA300_RS05230 | *purM* | phosphoribosylformylglycinamidine cyclo-ligase | | -1.22 |
| SAUSA300_RS05240 | *purH* | bifunctional purine biosynthesis protein | | -1.10 |
| SAUSA300_RS05245 | *purD* | phosphoribosylamine--glycine ligase | | -1.50 |
| SAUSA300_RS05275 | - | hypothetical protein | -2.07 | -2.28 |
| SAUSA300_RS05290 | *ptsH* | phosphocarrier protein | | -3.59 |
| SAUSA300_RS05295 | *ptsI* | phosphoenolpyruvate--protein phosphotransferase | -2.72 | -3.17 |
| SAUSA300_RS05355 | *pdhB* | pyruvate dehydrogenase E1 component subunit beta | -3.30 | -3.30 |
| SAUSA300_RS05360 | *pdhC* | 2-oxo acid dehydrogenase subunit E2 | | -1.99 |
| SAUSA300_RS05395 | *potD* | spermidine/putrescine ABC transporter substrate-binding protein | -1.19 | -1.51 |
| SAUSA300_RS05400 | - | hypothetical protein | | -1.24 |
| SAUSA300_RS05425 | - | hypothetical protein | | -1.01 |
| SAUSA300_RS05460 | *ctaA* | heme A synthase | -2.80 | -4.24 |
| SAUSA300_RS05465 | *ctaB* | protoheme IX farnesyltransferase | -1.65 | -3.74 |
| SAUSA300_RS05470 | *ctaM* | membrane protein | -1.36 | -2.43 |
| SAUSA300_RS05480 | - | hypothetical protein | | -1.53 |
| SAUSA300_RS05520 | - | DNA-binding protein | -2.90 | -2.20 |
| SAUSA300_RS05575 | - | hypothetical protein | | -2.62 |
| SAUSA300_RS05595 | *rnhC* | ribonuclease HIII | -1.53 | -1.37 |
| SAUSA300_RS05635 | *sdhC* | succinate dehydrogenase cytochrome B558 | -1.33 | -2.28 |
| SAUSA300_RS05640 | *sdhA* | succinate dehydrogenase flavoprotein subunit | | -1.74 |
| SAUSA300_RS05645 | *sdhB* | succinate dehydrogenase iron-sulfur subunit | | -1.30 |
| SAUSA300_RS05905 | *lspA* | lipoprotein signal peptidase | -3.59 | -3.63 |
| SAUSA300_RS05910 | - | RluA family pseudouridine synthase | -3.62 | -3.11 |
| SAUSA300_RS05915 | *pyrR* | bifunctional pyr operon transcriptional regulator/uracil phosphoribosyltransferase | -1.26 | -1.76 |
| SAUSA300_RS05920 | *pyrP* | uracil permease | -2.03 | -2.79 |
| SAUSA300_RS05925 | *pyrB* | aspartate carbamoyltransferase | | -1.56 |

| | | | | |
|---|---|---|---|---|
| SAUSA300_RS05930 | *pyrC* | dihydroorotase | -1.02 | -2.59 |
| SAUSA300_RS05935 | *carA* | carbamoyl-phosphate synthase small chain | | -1.33 |
| SAUSA300_RS05940 | *carB* | carbamoyl-phosphate synthase large chain | | -2.21 |
| SAUSA300_RS05945 | *pyrF* | orotidine-5'-phosphate decarboxylase | | -1.48 |
| SAUSA300_RS06000 | *def2* | peptide deformylase | -1.35 | -1.31 |
| SAUSA300_RS06010 | *sun* | 16S rRNA (cytosine(967)-C(5))-methyltransferase | -1.21 | -1.42 |
| SAUSA300_RS06015 | *rlmN* | 23S rRNA (adenine(2503)-C(2))-methyltransferase | -1.32 | -1.13 |
| SAUSA300_RS06020 | *stp1* | protein phosphatase | -2.98 | -1.85 |
| SAUSA300_RS06060 | *recG* | DNA helicase | -3.08 | -3.11 |
| SAUSA300_RS06095 | *rnc* | ribonuclease 3 | -1.10 | -2.01 |
| SAUSA300_RS06100 | *smc* | chromosome segregation protein | -2.61 | -2.58 |
| SAUSA300_RS06110 | - | DNA-binding protein | | -3.22 |
| SAUSA300_RS06135 | *rplS* | 50S ribosomal protein L19 | -2.39 | |
| SAUSA300_RS06160 | *sucC* | succinyl-CoA ligase subunit beta | | -2.57 |
| SAUSA300_RS06165 | *sucD* | succinyl-CoA ligase subunit alpha | -1.25 | |
| SAUSA300_RS06195 | *xerC* | tyrosine recombinase | | -2.61 |
| SAUSA300_RS06210 | *codY* | GTP-sensing pleiotropic transcriptional regulator | -2.35 | -3.02 |
| SAUSA300_RS06270 | *rimP* | ribosome maturation factor | -4.96 | -5.03 |
| SAUSA300_RS06285 | *rplGA* | hypothetical protein | -2.99 | -2.97 |
| SAUSA300_RS06295 | *rbfA* | ribosome-binding factor A | -2.78 | -2.46 |
| SAUSA300_RS06310 | *rpsO* | 30S ribosomal protein S15 | | -3.79 |
| SAUSA300_RS06315 | *pnpA* | polyribonucleotide nucleotidyltransferase | -3.09 | -4.08 |
| SAUSA300_RS06325 | *spoIIIE* | DNA translocase | -1.96 | -1.80 |
| SAUSA300_RS06340 | - | peptidase M16 | -1.39 | -1.32 |
| SAUSA300_RS06355 | - | transcriptional regulator | -2.98 | |
| SAUSA300_RS06375 | *rny* | ribonuclease Y | -4.01 | -4.65 |
| SAUSA300_RS06380 | - | hypothetical protein | | -1.19 |
| SAUSA300_RS06385 | - | metallophosphoesterase | -4.53 | -4.31 |
| SAUSA300_RS06390 | - | 2-oxoglutarate ferredoxin oxidoreductase subunit alpha | -1.81 | -1.32 |
| SAUSA300_RS06395 | - | 2-oxoacid ferredoxin oxidoreductase subunit beta | -1.61 | -1.50 |
| SAUSA300_RS06445 | *glpD* | aerobic glycerol-3-phosphate dehydrogenase | -2.22 | -3.39 |
| SAUSA300_RS06735 | - | hypothetical protein | | -4.69 |
| SAUSA300_RS06740 | - | membrane protein | | -2.16 |
| SAUSA300_RS06765 | *citB* | aconitate hydratase | | -1.91 |
| SAUSA300_RS06790 | *parC* | DNA topoisomerase 4 subunit A | -2.79 | -2.93 |
| SAUSA300_RS06830 | *msrR* | regulatory protein | -1.46 | -1.90 |
| SAUSA300_RS07035 | *cspA* | cold-shock protein | | -3.97 |
| SAUSA300_RS07050 | *acyP* | acylphosphatase | | -1.12 |

| | | | | |
|---|---|---|---|---|
| SAUSA300_RS07100 | *sucB* | dihydrolipoyllysine-residue succinyltransferase | | -2.50 |
| SAUSA300_RS07105 | *sucA* | 2-oxoglutarate dehydrogenase E1 component | | -1.57 |
| SAUSA300_RS07355 | *bshA* | N-acetyl-alpha-D-glucosaminyl L-malate synthase | -1.81 | -1.96 |
| SAUSA300_RS07360 | - | hypothetical protein | | -1.05 |
| SAUSA300_RS07390 | *aroB* | 3-dehydroquinate synthase | | -1.96 |
| SAUSA300_RS07405 | *ndk* | nucleoside-diphosphate kinase | | -1.53 |
| SAUSA300_RS07435 | *gpsA* | glycerol-3-phosphate dehydrogenase (NAD(P)(+)) | -1.08 | -1.16 |
| SAUSA300_RS07500 | *fer* | ferredoxin | -4.16 | -5.11 |
| SAUSA300_RS07820 | - | transcriptional regulator | | -1.44 |
| SAUSA300_RS07875 | *srrA* | DNA-binding response regulator | -1.65 | -3.00 |
| SAUSA300_RS07885 | *scpB* | SMC-Scp complex subunit | -2.93 | -5.24 |
| SAUSA300_RS07890 | *scpA* | segregation and condensation protein A | -3.99 | -4.38 |
| SAUSA300_RS07900 | *xerD* | tyrosine recombinase | -1.73 | -1.70 |
| SAUSA300_RS07990 | - | hypothetical protein | | -1.59 |
| SAUSA300_RS07995 | *bmfBB* | 2-oxoglutarate dehydrogenase E2 | -1.39 | -1.52 |
| SAUSA300_RS08000 | *bfmBAB* | alpha-ketoacid dehydrogenase subunit beta | -4.00 | -2.61 |
| SAUSA300_RS08005 | *bfmBAA* | 2-oxoisovalerate dehydrogenase subunit alpha | -2.88 | -4.69 |
| SAUSA300_RS08010 | *lpdA* | dihydrolipoyl dehydrogenase | -3.76 | -3.73 |
| SAUSA300_RS08015 | *recN* | DNA repair protein | -3.27 | -3.32 |
| SAUSA300_RS08020 | *argR* | arginine repressor | -1.22 | -2.00 |
| SAUSA300_RS08025 | *ispA* | geranyltranstransferase | -3.18 | -3.27 |
| SAUSA300_RS08030 | *xseB* | exodeoxyribonuclease 7 small subunit | | -1.11 |
| SAUSA300_RS08035 | *xseA* | exodeoxyribonuclease 7 large subunit | | -1.01 |
| SAUSA300_RS08040 | *nusB* | N utilization substance protein B | -2.43 | -3.67 |
| SAUSA300_RS08045 | - | Asp23/Gls24 family envelope stress response protein | -3.05 | -2.41 |
| SAUSA300_RS08180 | *aroK* | shikimate kinase | -1.05 | |
| SAUSA300_RS08240 | - | rhomboid family intramembrane serine protease | -2.31 | -2.29 |
| SAUSA300_RS08260 | *sodA* | superoxide dismutase | -3.47 | -4.31 |
| SAUSA300_RS08285 | *cshB* | DEAD/DEAH box family ATP-dependent RNA helicase | -1.39 | -4.37 |
| SAUSA300_RS08310 | - | phosphoenolpyruvate synthetase regulatory protein | -1.28 | -1.53 |
| SAUSA300_RS08325 | *recO* | DNA repair protein | | -4.36 |
| SAUSA300_RS08335 | *cdd* | cytidine deaminase | -1.59 | -1.24 |
| SAUSA300_RS08340 | *dgkA* | diacylglycerol kinase | -1.13 | -1.50 |
| SAUSA300_RS08350 | *phoH* | PhoH family protein | | -1.69 |
| SAUSA300_RS08370 | *rpsU* | 30S ribosomal protein S21 | -1.09 | -3.44 |
| SAUSA300_RS08405 | *hrcA* | HrcA family transcriptional regulator | -2.80 | -3.03 |
| SAUSA300_RS08420 | *lepA* | elongation factor 4 | -3.95 | -3.92 |
| SAUSA300_RS08460 | - | haloacid dehalogenase | | -1.18 |

| | | | | |
|---|---|---|---|---|
| SAUSA300_RS08475 | *aroE* | shikimate dehydrogenase | -1.17 | -1.39 |
| SAUSA300_RS08490 | *mtnN* | 5'-methylthioadenosine/S-adenosylhomocysteine nucleosidase | -4.62 | -4.42 |
| SAUSA300_RS08545 | *greA* | transcription elongation factor | -4.32 | -4.43 |
| SAUSA300_RS08550 | *udk* | uridine kinase | | -1.05 |
| SAUSA300_RS08570 | - | hypothetical protein | -1.67 | -2.72 |
| SAUSA300_RS08575 | *ruvX* | Holliday junction resolvase | -2.31 | -1.47 |
| SAUSA300_RS08580 | - | hypothetical protein | | -2.16 |
| SAUSA300_RS08590 | *recD2* | hypothetical protein | -2.59 | -2.45 |
| SAUSA300_RS08625 | *cymR* | Rrf2 family transcriptional regulator | -4.85 | -4.68 |
| SAUSA300_RS08665 | *relA* | bifunctional (p)ppGpp synthetase/hydrolase | | -1.15 |
| SAUSA300_RS08680 | *secDF* | protein translocase subunit | | -2.64 |
| SAUSA300_RS08700 | *ruvB* | Holliday junction branch migration DNA helicase | -1.69 | -2.90 |
| SAUSA300_RS08710 | *thrR* | hypothetical protein | -1.26 | |
| SAUSA300_RS08740 | *mreC* | rod shape-determining protein | | -1.70 |
| SAUSA300_RS08800 | *hemL* | glutamate-1-semialdehyde 2,1-aminomutase | | -1.49 |
| SAUSA300_RS08820 | *hemX* | cytochrome c assembly protein | -1.38 | |
| SAUSA300_RS08835 | *clpX* | ATP-dependent Clp protease ATP-binding subunit | | -2.02 |
| SAUSA300_RS08840 | *tig* | trigger factor | | -1.20 |
| SAUSA300_RS08855 | *mutT* | DNA mismatch repair protein | -2.20 | -2.06 |
| SAUSA300_RS08925 | *polA* | DNA polymerase I | -3.36 | -3.33 |
| SAUSA300_RS08950 | *citC* | isocitrate dehydrogenase (NADP(+)) | -1.23 | |
| SAUSA300_RS08955 | *citZ* | citrate synthase | -3.16 | -3.06 |
| SAUSA300_RS08980 | *pfkA* | ATP-dependent 6-phosphofructokinase | | -1.36 |
| SAUSA300_RS08995 | - | NAD-dependent malic enzyme 4 | -1.75 | -1.82 |
| SAUSA300_RS09040 | *uspA1* | universal stress protein | | -2.35 |
| SAUSA300_RS09045 | *ackA* | acetate kinase | | -1.06 |
| SAUSA300_RS09065 | *thiI* | tRNA 4-thiouridine(8) synthase | | -2.87 |
| SAUSA300_RS09185 | *ccpA* | catabolite control protein A | -1.03 | |
| SAUSA300_RS09190 | - | hypothetical protein | | -2.26 |
| SAUSA300_RS09195 | *aroA2* | bifunctional 3-deoxy-7-phosphoheptulonate synthase/chorismate mutase | -1.08 | |
| SAUSA300_RS09245 | - | peptidase | | -1.43 |
| SAUSA300_RS09430 | - | hypothetical protein | -1.10 | |
| SAUSA300_RS09470 | *pckA* | phosphoenolpyruvate carboxykinase (ATP) | | -2.13 |
| SAUSA300_RS09500 | *menE* | 2-succinylbenzoate-CoA ligase | -1.17 | |
| SAUSA300_RS09750 | *hemY* | protoporphyrinogen oxidase | -2.26 | -2.23 |
| SAUSA300_RS09760 | *hemE* | uroporphyrinogen decarboxylase | -3.09 | -4.25 |
| SAUSA300_RS09765 | - | hypothetical protein | -3.07 | -4.91 |
| SAUSA300_RS09775 | - | multidrug ABC transporter ATP-binding protein | -1.24 | -1.56 |

| | | | | |
|---|---|---|---|---|
| SAUSA300_RS09805 | *cbf1* | 3'-5' exoribonuclease | | -5.63 |
| SAUSA300_RS09810 | - | DNA double-strand break repair Rad50 ATPase | | -1.19 |
| SAUSA300_RS09815 | - | DNA repair exonuclease | | -3.78 |
| SAUSA300_RS10060 | *perR* | transcriptional repressor | -1.16 | -4.63 |
| SAUSA300_RS10085 | - | hypothetical protein | | -2.38 |
| SAUSA300_RS10135 | *sgtB* | monofunctional glycosyltransferase | | -1.85 |
| SAUSA300_RS10185 | *vraR* | DNA-binding response regulator | | -1.95 |
| SAUSA300_RS10190 | *vraS* | two-component sensor histidine kinase | -3.50 | -4.09 |
| SAUSA300_RS10195 | *vraT* | transporter | -2.63 | -2.92 |
| SAUSA300_RS10315 | *ligA* | DNA ligase (NAD(+)) | -3.14 | -2.67 |
| SAUSA300_RS10335 | *purB* | adenylosuccinate lyase | -1.44 | -2.26 |
| SAUSA300_RS10415 | - | hypothetical protein | -1.38 | -2.74 |
| SAUSA300_RS10455 | *pmtD* | membrane protein | -2.09 | |
| SAUSA300_RS10460 | *pmtC* | ABC transporter ATP-binding protein | -1.23 | |
| SAUSA300_RS10465 | *pmtB* | ABC-2 transporter family protein | -1.53 | -2.15 |
| SAUSA300_RS10470 | *pmtA* | ABC transporter ATP-binding protein | | -3.23 |
| SAUSA300_RS10810 | - | XRE family transcriptional regulator | -1.98 | -3.99 |
| SAUSA300_RS10900 | *groEL* | molecular chaperone | -2.41 | -2.38 |
| SAUSA300_RS10990 | *rex* | transcriptional regulator | -3.05 | -3.02 |
| SAUSA300_RS11165 | - | membrane protein | -2.45 | -2.42 |
| SAUSA300_RS11210 | *cshA* | DEAD/DEAH box family ATP-dependent RNA helicase | -3.10 | -3.07 |
| SAUSA300_RS11335 | *atpD* | ATP synthase subunit beta | -3.89 | -3.71 |
| SAUSA300_RS11340 | *atpG* | ATP synthase subunit gamma | -3.58 | -3.55 |
| SAUSA300_RS11345 | *atpA* | ATP synthase subunit alpha | -5.59 | -6.20 |
| SAUSA300_RS11350 | *atpH* | ATP synthase subunit delta | -4.84 | -2.32 |
| SAUSA300_RS11365 | *atpB* | ATP synthase subunit A | | -1.84 |
| SAUSA300_RS11370 | - | ATP synthase | | -1.11 |
| SAUSA300_RS11405 | *prmC* | protein-(glutamine-N5) methyltransferase, release factor-specific | -3.70 | -2.67 |
| SAUSA300_RS11415 | *tdk* | thymidine kinase | | -1.32 |
| SAUSA300_RS11425 | *rho* | transcription termination factor | | -1.09 |
| SAUSA300_RS11435 | *qsrR* | transcriptional regulator | | -4.64 |
| SAUSA300_RS11445 | *fbaA* | fructose-bisphosphate aldolase | | -1.74 |
| SAUSA300_RS11460 | *rpoE* | DNA-directed RNA polymerase subunit delta | | -1.54 |
| SAUSA300_RS11545 | *manA* | mannose-6-phosphate isomerase | | -1.89 |
| SAUSA300_RS11570 | - | hypothetical protein | -4.42 | -4.09 |
| SAUSA300_RS11705 | *salA* | chromosome partitioning protein | -3.47 | -4.35 |
| SAUSA300_RS11715 | *sepA* | multidrug resistance efflux pump | -4.22 | -4.11 |
| SAUSA300_RS11740 | - | hypothetical protein | -2.67 | -4.84 |

| | | | | |
|---|---|---|---|---|
| SAUSA300_RS12000 | *ecfT* | energy-coupling factor transporter protein | -2.40 | |
| SAUSA300_RS12005 | *cbiO2* | energy-coupling factor transporter ATPase | -1.97 | |
| SAUSA300_RS12010 | *cbiO* | energy-coupling factor transporter ATPase | | -1.29 |
| SAUSA300_RS12200 | - | hypothetical protein | | -1.15 |
| SAUSA300_RS12205 | - | hypothetical protein | -1.53 | |
| SAUSA300_RS12215 | *femX* | lipid II:glycine glycyltransferase | | -1.15 |
| SAUSA300_RS12585 | *hutI* | imidazolonepropionase | -1.03 | -2.12 |
| SAUSA300_RS12610 | *lyrA* | lysostaphin resistance protein A | | -1.13 |
| SAUSA300_RS12615 | *rpiA* | ribose-5-phosphate isomerase | | -3.82 |
| SAUSA300_RS12770 | - | DUF3021 domain-containing protein | -1.08 | -1.07 |
| SAUSA300_RS12775 | *mqo* | malate:quinone oxidoreductase | -5.32 | -6.41 |
| SAUSA300_RS12820 | - | membrane protein | | -1.25 |
| SAUSA300_RS13250 | - | hypothetical protein | | -1.66 |
| SAUSA300_RS13475 | *pgcA* | phosphoglucomutase | | -1.67 |
| SAUSA300_RS13520 | *gtaB* | UTP--glucose-1-phosphate uridylyltransferase | -1.59 | -2.46 |
| SAUSA300_RS13615 | *fbp* | fructose 1,6-bisphosphatase | | -1.34 |
| SAUSA300_RS13790 | *mvaA* | hydroxymethylglutaryl-CoA reductase, degradative | -3.24 | -3.21 |
| SAUSA300_RS13795 | *mvaS* | hydroxymethylglutaryl-CoA synthase | -3.93 | -3.87 |
| SAUSA300_RS13915 | *isaA* | transglycosylase IsaA | -2.04 | -4.22 |
| SAUSA300_RS14015 | *pyrD* | dihydroorotate dehydrogenase (quinone) | | -5.58 |
| SAUSA300_RS14675 | *noc* | nucleoid occlusion protein | -3.20 | -3.57 |
| SAUSA300_RS14685 | *mnmG* | tRNA uridine-5-carboxymethylaminomethyl(34) synthesis enzyme | | -1.12 |
| SAUSA300_RS14690 | *mnmE* | tRNA uridine-5-carboxymethylaminomethyl(34) synthesis GTPase | -1.32 | -2.93 |
| SAUSA300_RS15990 | - | hypothetical protein | | -2.00 |

| Genes with significant increase in fitness during biofilm growth due to transposon insertions | | | | |
|---|---|---|---|---|
| | | | Log$_2$ Fold Change (vs inoculum) | |
| Gene_ID | Gene Name | Gene Product | 48-hour Biofilm | 96-hour Biofilm |
| SAUSA300_RS00590 | *sarS* | transcriptional regulator | | 1.11 |
| SAUSA300_RS00715 | - | GntR family transcriptional regulator | | 1.01 |
| SAUSA300_RS00745 | *phnE1* | phosphonate ABC transporter, permease protein | | 1.01 |
| SAUSA300_RS00960 | - | hypothetical protein | | 1.50 |
| SAUSA300_RS01520 | *esaC* | type VII secretion substrate | | 1.53 |
| SAUSA300_RS01545 | *esaG* | TIGR01741 family protein | | 1.02 |
| SAUSA300_RS01605 | - | TIGR01741 family protein | | 1.06 |

| | | | | |
|---|---|---|---|---|
| SAUSA300_RS02230 | *lpl6* | tandem-type lipoprotein | | 1.10 |
| SAUSA300_RS02535 | *ipk* | 4-diphosphocytidyl-2C-methyl-D-erythritol kinase | | 1.37 |
| SAUSA300_RS02540 | *purR* | pur operon repressor | 1.16 | 1.85 |
| SAUSA300_RS02545 | *yabJ* | RidA family protein | 1.63 | 2.00 |
| SAUSA300_RS03455 | *graX* | NAD(P)-dependent oxidoreductase | 1.25 | 2.00 |
| SAUSA300_RS03460 | *graR* | DNA-binding response regulator | 1.56 | 2.12 |
| SAUSA300_RS03465 | *graS* | sensor histidine kinase | 1.70 | 2.44 |
| SAUSA300_RS03470 | *vraF* | ABC transporter ATP-binding protein | 1.52 | 2.01 |
| SAUSA300_RS03475 | *vraG* | ABC transporter permease | 1.68 | 2.49 |
| SAUSA300_RS03605 | *mgrA* | MarR family transcriptional regulator | 1.00 | 1.23 |
| SAUSA300_RS04105 | - | hypothetical protein | | 1.03 |
| SAUSA300_RS04240 | - | N-acetyltransferase | | 1.55 |
| SAUSA300_RS04740 | - | 2-isopropylmalate synthase | | 1.04 |
| SAUSA300_RS05155 | *fmtA* | teichoic acid D-Ala esterase FmtA | 1.80 | 2.49 |
| SAUSA300_RS05780 | - | hypothetical protein | | 1.17 |
| SAUSA300_RS06200 | *hslV* | HslU--HslV peptidase proteolytic subunit | | 2.02 |
| SAUSA300_RS06820 | *mprF* | phosphatidylglycerol lysyltransferase | | 1.04 |
| SAUSA300_RS06855 | *trpE* | anthranilate synthase component I | | 1.03 |
| SAUSA300_RS07850 | - | hypothetical protein | | 1.34 |
| SAUSA300_RS10445 | - | membrane protein | 1.25 | 1.48 |
| SAUSA300_RS10935 | *agrB* | accessory gene regulator | 2.81 | 3.09 |
| SAUSA300_RS10940 | *agrD* | cyclic lactone autoinducer peptide | 2.28 | 2.59 |
| SAUSA300_RS10945 | *agrC* | ATP-binding protein | 2.53 | 2.89 |
| SAUSA300_RS10950 | *agrA* | DNA-binding response regulator | 2.86 | 3.15 |
| SAUSA300_RS10970 | *scrR* | LacI family transcriptional regulator | | 1.24 |
| SAUSA300_RS11120 | *sigB* | RNA polymerase sigma factor | 1.50 | 1.57 |
| SAUSA300_RS11125 | *rsbW* | anti-sigma B factor | 1.64 | 1.89 |
| SAUSA300_RS11130 | *rsbV* | anti-sigma B factor antagonist | 1.76 | 1.39 |
| SAUSA300_RS11135 | *rsbU* | serine phosphatase | 1.64 | 1.95 |
| SAUSA300_RS11145 | *mazF* | type II toxin-antitoxin system PemK/MazF family toxin | 1.48 | 1.87 |
| SAUSA300_RS11150 | *mazE* | type II toxin-antitoxin system antitoxin MazE | 1.60 | |
| SAUSA300_RS11380 | *upp* | uracil phosphoribosyltransferase | | 1.66 |
| SAUSA300_RS11385 | *glyA* | serine hydroxymethyltransferase | | 1.31 |
| SAUSA300_RS11390 | - | TIGR01440 family protein | | 1.16 |
| SAUSA300_RS11635 | *ybbR* | YbbR-like domain-containing protein | | 1.48 |
| SAUSA300_RS13415 | - | single-stranded DNA-binding protein | | 1.12 |
| SAUSA300_RS13785 | - | hypothetical protein | | 1.13 |
| SAUSA300_RS14125 | - | sterile alpha motif-like domain-containing protein | | 1.63 |

| | | | | |
|---|---|---|---|---|
| SAUSA300_RS14270 | *clfB* | clumping factor B | | 1.07 |
